# Global analysis of the heparin-enriched plasma proteome captures matrisome-associated proteins in Alzheimer’s disease

**DOI:** 10.1101/2023.11.06.565824

**Authors:** Qi Guo, Lingyan Ping, Eric B. Dammer, Duc M. Duong, Luming Yin, Kaiming Xu, Ananth Shantaraman, Edward J. Fox, Erik C.B. Johnson, Blaine R. Roberts, James J. Lah, Allan I. Levey, Nicholas T. Seyfried

**Affiliations:** Department of Biochemistry, School of Medicine, Emory University, Atlanta, GA 30322, USA; Department of Neurology, Emory University School of Medicine, Atlanta, GA, USA; Goizueta Alzheimer’s Disease Research Center, Emory University School of Medicine, Atlanta, GA, USA

**Keywords:** label free quantification, tandem mass tag labeling, mass spectrometry, Alzheimer’s disease, human plasma, heparin binding proteins

## Abstract

Matrisome-associated heparin binding proteins (HBPs) with roles in extracellular matrix assembly are strongly correlated to β-amyloid (Aβ) and tau pathology in Alzheimer’s disease (AD) brain and cerebrospinal fluid (CSF). However, it remains challenging to detect these proteins in plasma using standard mass spectrometry (MS)-based proteomic approaches. Here we utilized heparin affinity chromatography for the capture and enrichment of HBPs in plasma from healthy control and individuals with AD. This method was highly reproducible and effectively enriched well-known HBPs like APOE and thrombin, while also efficiently depleting high-abundance proteins such as albumin. To increase the depth of our analysis of the heparin-enriched plasma proteome and compare differences in disease we applied off-line fractionation and tandem mass tag mass spectrometry (TMT-MS) to compare the proteomic profiles between AD and control individuals across two datasets (*n* = 121 total samples). This led to the identification of 2865 proteins, spanning 10 orders of magnitude in protein abundance within the plasma. Notably, HBPs were some of the most increased proteins in AD plasma compared to controls. This included members of the matrisome-associated module in brain, SMOC1, SMOC2, SPON1, MDK, OLFML3, FRZB, GPNMB and the ɛ4 isoform of APOE. Heparin-enriched plasma proteins also exhibited strong correlations to conventional AD biomarkers including CSF Aβ, total tau (tTau), and phosphorylated tau (pTau) as well as plasma pTau supporting their role as potential surrogate markers of underlying brain pathology. Utilizing a consensus AD brain protein co-expression network, we assessed relationship between the plasma and brain proteomes and observed that specific plasma proteins exhibited consistent direction of change in both brain and plasma, whereas others displayed divergent changes, further highlighting the complex interplay between the two compartments. In summary, these findings provide support for the integration of a heparin enrichment method with MS-based proteomics for identifying a wide spectrum of plasma biomarkers that mirror pathological changes in the AD brain.

## Introduction

Alzheimer’s disease (AD) is an age driven neurodegenerative disease characterized by the accumulation of two core pathologies, amyloid beta (Aβ) plaques and phosphorylated tau neurofibrillary tangles (NFTs) in the brain, ultimately leading to dementia ^1-3^. Notably, both pathologies accumulate over a decade prior to clinical symptoms in a latent preclinical phase of disease, which provides an opportunity for early detection using biomarkers ^4^. To this end, progress towards early accurate diagnosis and effective treatments for AD has been mainly focused on the two hallmark pathologies with recent advances in developing assays for Aβ, tau and phosphorylated tau (pTau) species in cerebrospinal fluid (CSF) and plasma ^5-7^. Specifically, phosphorylated tau on threonine-181 (pTau181) and on threonine-217 (pTau217) have recently emerged as promising plasma biomarkers for AD, reflecting Aβ deposition as well as subsequent tau deposition in the brain ^5-7^. However, it has been demonstrated that combining multiple proteins in CSF enhances the accuracy and discriminative capability of both pTau and Aβ for dementia ^8^, which could apply to plasma as well.

Emerging evidence has suggested that Aβ and tau, represent only a fraction of the complex and heterogeneous biology of AD ^9, 10^. For example, large-scale bulk RNA-seq and proteomics studies link AD to diverse biological mechanisms beyond Aβ and tau involving various biochemical pathways and cell types in brain ^11-13^. These studies revealed pathophysiological mechanisms such as synapse loss and neuroinflammation linked to immune, vascular, metabolic and extracellular matrix (ECM) dysfunction. Furthermore, integrated analyses across brain and biofluids demonstrated a significant overlap between brain network changes and the CSF proteome in AD, enabling early disease prediction even in the preclinical phase of AD ^12, 14-16^. For example, CSF proteomic measurements in autosomal dominant AD (ADAD) that overlap with brain protein co-expression modules were recently used to define the evolution of AD pathology over a timescale spanning six decades ^17^. SMOC1 and SPON1, ECM proteins associated with Aβ plaques, showed elevated levels in CSF almost three decades before symptom onset. Subsequent alterations were observed in synaptic, metabolic, axonal, inflammatory proteins, and, finally, reductions in neurosecretory proteins ^17^. Similar trends in these biomarkers were observed in late-onset AD (LOAD) where a targeted CSF proteomic panel reflecting diverse brain-based pathophysiology enhanced the ability of Aβ, tau, and pTau in predicting clinical diagnosis, FDG PET, hippocampal volume, and measures of cognitive severity ^8^. Notably, *in vivo* measurements of fibrillary amyloid plaques in the brain of AD patients using the PET ligand florbetapir (AV45) was most strongly associated with SMOC1, further supporting its role as an important surrogate marker of underlying Aβ pathology in brain ^8^.

In a consensus human brain proteome network, both SMOC1 and SPON1 are hub proteins within module 42 (M42), which was assigned the term ‘matrisome’, given the collection of ECM-associated proteins and strong enrichment of glycosaminoglycan-binding proteins ^11, 18^. This module also contained several additional proteins that have previously been identified to be correlated with or directly bind to Aβ ^11, 19, 20^. This includes amyloid precursor protein (APP), a proteomic surrogate for Aβ deposition in brain ^21^, and apolipoprotein E (APOE) the protein product of the AD genetic risk factor *APOE ^22^*. Interestingly, levels of the M42 matrisome module were increased in individuals carrying the APOE ε4 allele, the strongest genetic risk factor for late-onset AD ^11^. Furthermore, most M42 proteins are heparan sulfate (HS) or heparin binding proteins (HBPs) including SMOC1, SPON1, MDK, and APOE among others ^23-26^. Notably, heparin and HS accelerate the formation of Aβ fibrils ^27-29^ and matrisome signaling has been associated with APOE ε4 in mixed cortical cultures ^30^. Taken together, these discoveries indicate that specific HBPs within M42, such as SMOC1 and SPON1, exhibiting associations with amyloid plaques, are linked to the APOE ε4 allele, and function as early indicators of AD risk in CSF. Hence, if readily detectable in plasma, members of M42 hold significant potential as biomarkers for amyloid pathology in AD.

Although SMOC1 and SPON1 have been reported to change in AD plasma using antibody- or aptamer-based proteomic technologies ^31^, members of M42 have been extremely difficult to identify and quantify using mass spectrometry (MS)-based proteomic approaches in plasma. Much like CSF, human plasma is characterized by a large dynamic range of protein abundance, estimated at 12–13 orders of magnitude ^32^, in which albumin and other highly abundant proteins can prevent the detection of proteins of interest. However, the concentration of albumin in plasma (∼640 μM) is about 200-fold higher than in CSF (∼3 μM), which means fractionation methods such as the immune-depletion of albumin and other highly abundant proteins are typically required to enhance the depth ^33, 34^. However, our attempts using these approaches have only partially enriched members of M42 in plasma ^31^. Therefore, given the shared heparin binding properties of M42 members, we aimed to capture and quantify M42 matrisome members from plasma using heparin affinity chromatography followed by MS-based proteomic analysis to enhance the coverage and quantification of these proteins and assess their changes in AD.

Here we describe a heparin affinity chromatography approach to capture and enrich HBPs in human plasma. Using this approach, we demonstrate that heparin affinity chromatography is effective not only in enriching known HBPs, APOE and thrombin, but also in depleting high-abundance proteins like albumin. To enhance the depth and throughput of the heparin-enriched plasma proteome, we utilized tandem mass tag mass spectrometry (TMT-MS) following high-pH off-line fractionation to determine differences in the proteome between AD and control individuals in replicate datasets (*n* = 121 samples total). Collectively we identified 2865 proteins across the two sets of samples spanning 10 orders of magnitude in protein abundance in plasma. We further show that members of M42 including SMOC1, SMOC2, SPON1, MDK, OLFML3, FRZB, GPNMB and APOE4 were significantly increased in AD plasma and correlated to CSF levels of amyloid, tau and pTau as well as plasma pTau suggesting that these proteins are related to AD pathophysiology in both brain and plasma. Finally, we leveraged the consensus brain protein co-expression network and examined the relationship between plasma and brain proteomes. Certain plasma proteins within the network modules showed consistent increases or decreases in both the AD brain and plasma, while others displayed a divergent change. In summary, these findings provide strong support for the integration of a heparin enrichment method with MS-based proteomic analysis for identifying a wide spectrum of plasma biomarkers that mirror pathological changes in the AD brain.

## Methods

### Materials

Primary antibodies used included a mouse monoclonal anti-prothrombin antibody (Catalog No. ab17199, Abcam) and a goat polyclonal anti-ApoE antibody (Catalog No. K74180B, Meridian Life Science). Secondary antibodies used were conjugated with either Alexa Fluor 680 (Invitrogen) or IRDye800 (Rockland) fluorophores for enhanced detection and visualization. Heparin-sepharose (Cytiva, lot#17099801 for Set 1 and lot#17099803 for Set 2) was used to enrich HBPs from plasma samples.

### Plasma and CSF Samples

All participants providing plasma and CSF samples gave their informed consent following the protocols approved by the Institutional Review Board at Emory University. Comprehensive cognitive assessments, including the Montreal Cognitive Assessment (MoCA), were administered to all patients as part of their evaluation at the Emory Cognitive Neurology Clinic, the Emory Goizueta Alzheimer’s Disease Research Center (ADRC), and related research initiatives such as the Emory Healthy Brain Study (EHBS). Diagnostic data were sourced from the ADRC and the Emory Cognitive Neurology Program. Plasma and CSF samples were collected from participants on the same day using standard procedures. These samples were processed and stored in accordance with the 2014 ADC/NIA best practices guidelines. For participants recruited through the Emory Cognitive Neurology Clinic, CSF samples were sent to Athena Diagnostics and assayed for CSF AD biomarkers, including Aβ^1-42^, tTau, and pTau181, utilizing the INNOTEST assay platform. CSF samples collected from research participants in the ADRC and EHBS were assayed using the INNO-BIA AlzBio3 Luminex assay. To analyze plasma pTau181 concentrations, EDTA plasma samples were prepared according to manufacturer’s instructions from the pTau181 kit v2 (Quanterix Billerica, Massachusetts, USA). Samples were run in a single batch. Plasma was thawed at room temperature for 45 minutes and then centrifuged at 5000×g for 10 minutes. The plasma samples were then diluted four times and measured on the Simoa HDX platform. Mean intra-assay coefficients of variation (CV) were below 10%. In total, a pooled plasma sample and two sets of individual plasma samples were used in the study. Set 1 comprised plasma samples from 18 cognitively normal healthy controls and 18 individuals with mild cognitive impairment or AD, whereas Set 2 included plasma samples obtained from 36 cognitively normal control individuals and 49 individuals with mild cognitive impairment or AD. 13 out of these 121 samples are overlapping in the two sets and thus a total of 108 unique cases. For the (Set 1 + Set 2) meta-analysis, 12 non-overlapping cases that did not meet strict CSF biomarker criteria (tTau/Aβ^1-42^ ratio > 0.226 for AD) or MoCA cutoffs (AD < 24, Control > 26) ^35, 36^ at the time of lumbar puncture were removed, ending up with 109 samples (unique cases = 96). Further details about the cohorts are available in **Table 1 and Supplemental Table 4 and 10**.

**Table 1.**
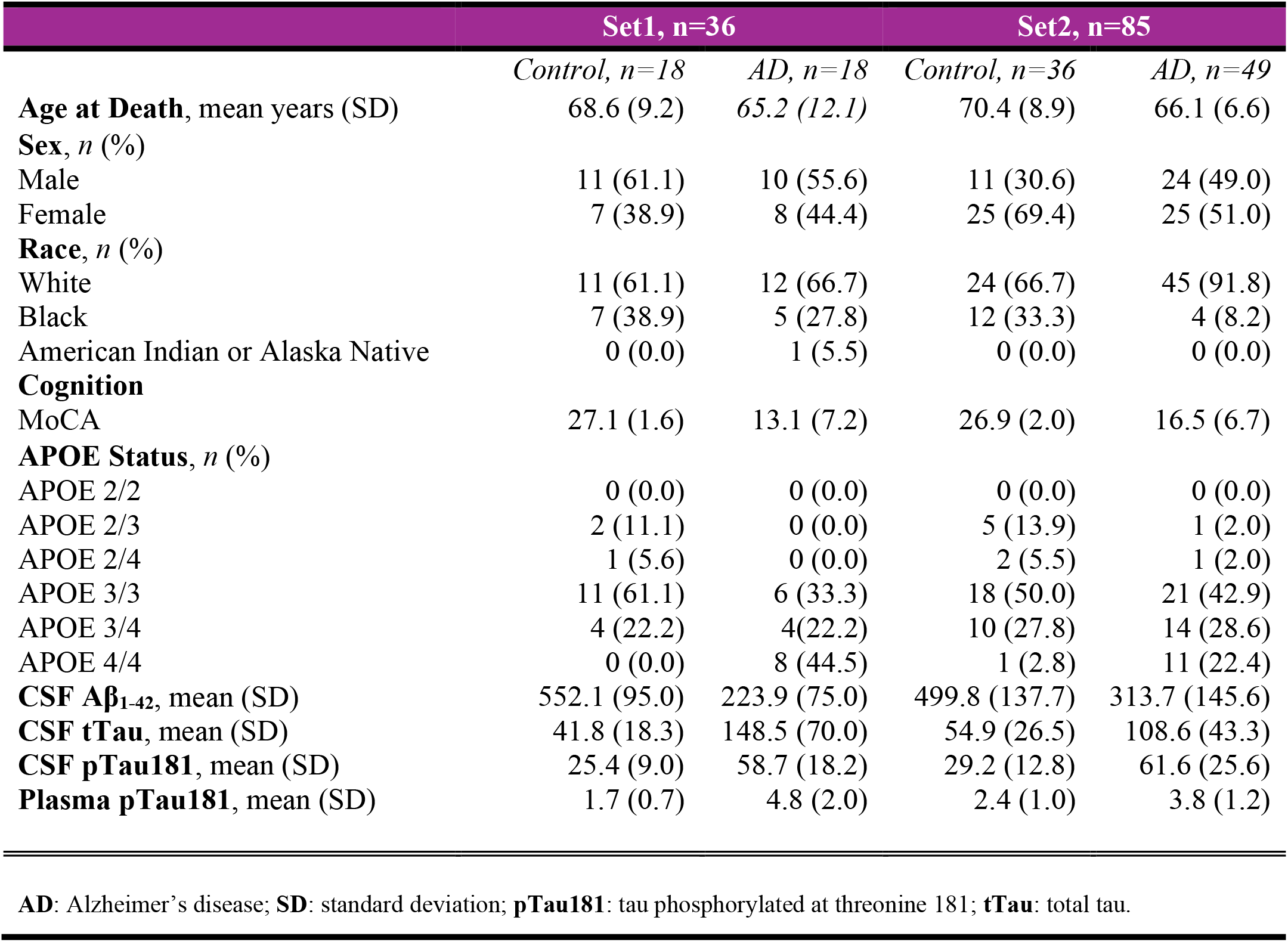
Characteristics of human subjects used in this study.

|  | Set1, n=36 |  | Set2, n=85 |  |
| --- | --- | --- | --- | --- |
|  | *Control, n=18* | *AD, n=18* | *Control, n=36* | *AD, n=49* |
| **Age at Death**, mean years (SD) | 68.6 (9.2) | 65.2 (12.1) | 70.4 (8.9) | 66.1 (6.6) |
| **Sex**, *n* (%) |  |  |  |  |
| Male | 11 (61.1) | 10 (55.6) | 11 (30.6) | 24 (49.0) |
| Female | 7 (38.9) | 8 (44.4) | 25 (69.4) | 25 (51.0) |
| **Race**, *n* (%) |  |  |  |  |
| White | 11 (61.1) | 12 (66.7) | 24 (66.7) | 45 (91.8) |
| Black | 7 (38.9) | 5 (27.8) | 12 (33.3) | 4 (8.2) |
| American Indian or Alaska Native | 0 (0.0) | 1 (5.5) | 0 (0.0) | 0 (0.0) |
| **Cognition** |  |  |  |  |
| MoCA | 27.1 (1.6) | 13.1 (7.2) | 26.9 (2.0) | 16.5 (6.7) |
| **APOE Status**, *n* (%) |  |  |  |  |
| APOE 2/2 | 0 (0.0) | 0 (0.0) | 0 (0.0) | 0 (0.0) |
| APOE 2/3 | 2 (11.1) | 0 (0.0) | 5 (13.9) | 1 (2.0) |
| APOE 2/4 | 1 (5.6) | 0 (0.0) | 2 (5.5) | 1 (2.0) |
| APOE 3/3 | 11 (61.1) | 6 (33.3) | 18 (50.0) | 21 (42.9) |
| APOE 3/4 | 4 (22.2) | 4 (22.2) | 10 (27.8) | 14 (28.6) |
| APOE 4/4 | 0 (0.0) | 8 (44.5) | 1 (2.8) | 11 (22.4) |
| **CSF Aβ<sub>1-42</sub>**, mean (SD) | 552.1 (95.0) | 223.9 (75.0) | 499.8 (137.7) | 313.7 (145.6) |
| **CSF tTau**, mean (SD) | 41.8 (18.3) | 148.5 (70.0) | 54.9 (26.5) | 108.6 (43.3) |
| **CSF pTau181**, mean (SD) | 25.4 (9.0) | 58.7 (18.2) | 29.2 (12.8) | 61.6 (25.6) |
| **Plasma pTau181**, mean (SD) | 1.7 (0.7) | 4.8 (2.0) | 2.4 (1.0) | 3.8 (1.2) |

**AD:** Alzheimer's disease; **SD:** standard deviation; **pTau181:** tau phosphorylated at threonine 181; **tTau:** total tau.

### Heparin binding protein enrichment from plasma

The heparin enrichment process was conducted in technical triplicates, using 40 µl of pooled neat human plasma per replicate. Initially, 100 µl of heparin-sepharose bead slurry (1:1 w/v) was prepared with 50 µl of beads for each replicate (*n* = 3) and binding buffer (50 mM sodium phosphate, pH 7.4). Each slurry was then washed twice with 1 ml of binding buffer. Subsequently, each replicate of neat plasma (40 µl) was diluted with 1 ml of binding buffer to generate diluted plasma (DP) as input and mixed with the heparin-sepharose beads prepared above. The incubation was performed at room temperature for 10 minutes coupled with rotation. Following the enrichment step, the beads were spun down at 500×g for 2 minutes and the supernatant was collected as the heparin-depleted flowthrough (Hp-depleted FT) fraction. The heparin-sepharose beads were then washed twice with 1 ml of binding buffer and resuspended in 1 ml of binding buffer (Hp-enriched fraction). The Hp-enriched fraction was then split into 650 µl for digestion and 350 µl for SDS-PAGE and western blotting. The supernatant was removed before further processing. For Set 1 (*n* = 36) and Set 2 (*n* = 85) samples, a global pooled standard (GPS) was prepared for each set as internal control for enrichment by pooling equal amount of each sample within each set. A nearly identical protocol was followed for subsequent enrichment, using 50 µl of beads (Set 1) and 200 µl of beads (Set 2) correspondingly for each set.

### Gel electrophoresis and western blot analysis

Western blotting and Coomassie Blue staining were performed on all three fractions from pooled plasma sample (DP input, *n* = 3; Hp-depleted FT, *n* = 3; Hp-enriched fraction, *n* = 3) as previously described ^37-39^. For Coomassie Blue staining, samples were boiled with 4x Laemmli sample buffer, and an equal volume to 0.2 µl of the neat plasma was loaded from DP inputs and FT fractions onto an SDS-PAGE gel (Invitrogen). To enhance protein visualization, the Hp-enriched fractions were loaded at a 5-fold higher amount (equal to 1 µl of the neat plasma) than the input and FT. For the western blotting, an equal volume to 0.25 µl of the neat plasma was loaded from all three fractions. The gels were either stained with Coomassie Blue G250 overnight or semi-dry transferred to nitrocellulose membranes (Invitrogen) using the iBlot2 system (Life Technologies). Subsequently, the membranes were blocked with casein blocking buffer (Sigma B6429) for 30 minutes at room temperature. They were then probed with two primary antibodies (mouse monoclonal anti-prothrombin and goat polyclonal anti-APOE) at a 1:1000 dilution overnight at 4 °C. On the following day, the membranes were rinsed and incubated with secondary antibodies conjugated to the Alexa Fluor 680 fluorophore (Invitrogen) at a 1:10000 dilution for one hour at room temperature. After another round of rinsing, the membranes were once again incubated with secondary antibodies conjugated to a second IRDye800 fluorophore at a 1:10000 dilution for one hour at room temperature. For Set 1 samples (*n* = 36), western blotting was performed on each of the sample as well as GPS as described above using an equal volume to 0.25 µl of the neat plasma from all three fractions. For Set 2 samples (*n* = 85), western blotting was performed on only the GPS in triplicates with an equal volume to 0.25 µl of the neat plasma from all three fractions.

### Heparin-enriched plasma protein digestion

Sample digestion was carried out on all three fractions for pooled plasma sample (DP input, *n* = 3; Hp-depleted FT, *n* = 3; Hp-enriched fraction, *n* = 3), and only Hp-enriched fraction for Set 1 (*n* = 36) and Set 2 (*n* = 85) samples. As previously described ^40^, equal volume (25 µl) of DP input and FT fractions from pooled plasma sample was reduced and alkylated with 2.5 µl of 0.4 M chloroacetamide (CAA) and 0.5 µl of 0.5 M tris-2(-carboxyethyl)-phosphine (TCEP, ThermoFisher) in a 90 °C water bath for 10 minutes. Following this, water bath sonication was performed for 5 minutes. Samples were then cooled down to room temperature before incubating with 50 mAU of lysyl endopeptidase (LysC, Wako) in 28 µl of 8 M urea buffer (8 M urea, 10mM Tris, 100 mM Na^2^HPO^4^, pH 8.5) and kept at 26 °C overnight for LysC digestion. The following day, 10 µg of trypsin (Promega) was added with 168 µl of 50 mM ammonium carbonate and kept at 26 °C overnight for trypsin digestion. For the Hp-enriched fractions, digestion was performed on beads. The supernatant was removed before adding 40 µl of 0.4 M CAA and 8 µl of 0.5 M TCEP with 400 µl of 50 mM ammonium carbonate for reduction and alkylation. Following the same steps above, 25 mAU of LysC was used for LysC digestion at 37 °C with shaking at 1000 rpm. 10 µg of trypsin was then added the following day for overnight digestion. After trypsin digestion, the digested peptides were acidified to a final concentration of 1% (vol/vol) formic acid (FA) and 0.1% (vol/vol) triflouroacetic acid (TFA) and desalted with a 10 mg HLB column (Waters). Each column was activated with 1 ml of methanol, followed by equilibration with 2×1 ml of 0.1% (vol/vol) TFA. The samples were then loaded onto the column and washed with 2×1 ml of 0.1% (vol/vol) TFA. Elution was performed with 2 rounds of 500 μl of 50% (vol/vol) acetonitrile (ACN). Global internal standard (GIS) was prepared for Set 1 and Set 2 separately by pooling 100 µl of the elution from all samples within each set and divided into 900 µl per aliquot. The elution was dried to completeness via speed vacuum (Labconco).

### Label-free mass spectrometry and quantification

Dried peptides were reconstituted in peptide loading buffer (0.1% FA, 0.03% TFA, 1% ACN). Using an RSLCnano liquid chromatography (LC) system, approximately 1 µg of peptide was loaded onto an in-house made column (75 µm internal diameter and 50 cm length) packed with 1.9-μm ReproSil-Pur C18-AQ resin (Maisch, Germany) and eluted over a 120-minute gradient. Elution was performed at a rate of 300 nl/min with buffer B/buffer (A+B) ratio ranging from 1 to 99% (buffer A, 0.1% FA in water; buffer B, 0.1% FA in 80% ACN). Mass spectrometry was performed with a high-field asymmetric waveform ion mobility spectrometry (FAIMS) Pro-equipped Eclipse Orbitrap mass spectrometer (ThermoFisher) in positive ion mode using data-dependent acquisition (DDA) with 3 x 1-second top speed cycles and 3 compensation voltages (- 40, -60 and -80). Each compensation voltage (CV) top speed cycle consisted of one full MS scan with as many MS/MS events that could fit in the 1-second cycle time. Full MS scans were collected at a resolution of 120k [350 to 1500 mass/charge ratio (m/z) range, 4 × 10^-5^ automatic gain control (AGC) target, and 50-ms maximum ion injection time]. All higher-energy collision-induced dissociation (HCD) MS/MS spectra were acquired in the ion trap (1.6 m/z isolation width, 35% collision energy, 1×10^-4^ AGC target, and 35-ms maximum ion time). Dynamic exclusion was set to exclude previously sequenced peaks for 60 s within a 10-ppm (parts per million) isolation window. Only precursor ions with charge states between 2 and 7 were selected for fragmentation.

### Database search parameters for label-free quantification of samples

The raw files of all three fractions from pooled plasma sample (*n* = 9) were searched using FragPipe (FP, version 20.0). The FP pipeline for label-free quantification (LFQ) relies on MSFragger ^41, 42^ (version 3.8) for peptide identification. The peptide search was done against all canonical human proteins downloaded from Uniprot (20,402 total sequencese; accessed 02/11/2019), as well as 51 common contaminants, and all 20453 reverse sequences (decoys). The prescribed LFQ-MBR workflow in FP was used with parameters specified as follows: precursor mass tolerance was -20 to 20 ppm, fragment mass tolerance of 0.7 Da, mass calibration and parameter optimization were selected, and isotope error was set to 0/1/2. Cleavage type was set to semi-enzymatic. Enzyme specificity was set to strict-trypsin and up to two missing trypsin cleavages were allowed. Peptide length was allowed to range from 7 to 50 and peptide mass from 200 to 5,000 Da. Variable modifications that were allowed in our search included: oxidation on methionine, and N-terminal acetylation on protein. Static modifications included: carbamidomethylation on cysteine. Peptide-spectrum match (PSM) were validated using Percolator ^43^. The false discovery rate (FDR) threshold was set to 1% using Philosopher ^44^ (version 5.0.0). The peptide and UniprotID-identified protein abundances were quantified using IonQuant ^45^ (version 1.9.8) for downstream analysis.

### Data filtration, normalization and imputation for LFQ

To enable protein overlap analysis across the three fractions and conduct gene ontology (GO) analysis for proteins within each fraction, proteins that were detected in at least 2 out of 3 replicates within each fraction were selected. Before performing differential abundance analysis, the data from all nine samples were merged and protein levels were first scaled by dividing each protein intensity by intensity sum of all proteins in each sample followed by multiplying by the maximum protein intensity sum across all nine samples. Instances where the intensity was ‘0’ were treated as ‘missing values’. Subsequently, Perseus-style imputation was applied to these missing values for further analysis ^46^.

### Tandem mass tag (TMT) labeling of peptides

TMT labeling of digested Set 1 and Set 2 samples was performed as previously described ^40^. The discovery dataset (Set 1) was comprised of 36 individual samples and 3 GIS randomized based on age, sex and diagnosis into three batches and the replication dataset (Set 2) was comprised of 85 individual samples and 5 GIS randomized into five batches. **Supplemental Table 4 and 10** provides the sample-to-batch arrangement for each of these samples. These samples were then labeled using TMTpro kits (Thermo Fisher Scientific, A44520, Lot number: VH3111511 for Set 1; UK297033 for Set 2 with XB338618 for channels 134C and 135). First, each peptide digest was resuspended in 75 μl of 100 mM triethylammonium bicarbonate (TEAB) buffer, and 5 mg of TMT reagent was dissolved into 200 μl of anhydrous acetonitrile (ACN). After that, 15 μl of TMT reagent solution was subsequently added to the resuspended peptide digest and incubated for 1 hour at room temperature. Following that, the reaction was quenched with 4 μl of 5% hydroxylamine (Thermo Fisher Scientific, 90115) for 15 minutes. Then, the peptide solutions were combined according to the batch arrangement. Finally, each TMT batch was desalted with 60 mg HLB columns (Waters) and dried via speed vacuum (Labconco).

### High-pH off-line peptide fractionation

TMT labeled samples were prepared for high-pH off-line fractionation as previously described ^40^ with slight modifications. Briefly, dried samples from above were re-suspended in high-pH loading buffer (0.07% vol/vol NH^4^OH, 0.045% vol/vol FA, 2% vol/vol ACN) and loaded onto a Waters BEH column (2.1 mm x 150 mm with 1.7 µm particles). A Vanquish UPLC system (Thermo Fisher Scientific) was used to carry out the fractionation. Solvent A consisted of 0.0175% (vol/vol) NH^4^OH, 0.01125% (vol/vol) FA, and 2% (vol/vol) ACN; solvent B consisted of 0.0175% (vol/vol) NH^4^OH, 0.01125% (vol/vol) FA, and 90% (vol/vol) ACN. The sample elution was performed over a 25-minute gradient (0 to 50% solvent B) with a flow rate of 0.6 ml/min. A total of 96 individual equal volume fractions were collected across the gradient per TMT batch and dried via speed vacuum (Labconco).

### Mass spectrometry analysis for TMT labeled samples

An equal volume of each high-pH peptide fraction was initially resuspended in loading buffer (0.1% FA, 0.03% TFA, and 1% ACN) and was loaded and separated using an EASY-nanoLC system on a self-made 15-cm-long, 150-μM internal diameter (ID) fused silica column packed with 1.9-μm ReproSil-Pur C18-AQ resin from Maisch, Germany. Elution was carried out over a 20-minute gradient at a flow rate of 1200 nL/min, with buffer B/buffer (A+B) ratio ranging from 1% to 99%. For Set 1, mass spectrometry was performed on a high-field asymmetric waveform ion mobility spectrometry (FAIMS) Pro-equipped Orbitrap Lumos (Thermo) in positive ion mode. DDA was used with 2 x 1-second top speed cycles and 2 compensation voltages (-45 and -65). Each top speed cycle included one full MS scan with as many MS/MS events as possible within the 1-second cycle time. Full MS scans were acquired at a resolution of 120,000 over a m/z range of 410 to 1600, with an AGC target of 4 × 10^-5^ and a maximum ion injection time of 50 ms. All HCD MS/MS spectra were obtained at a resolution of 50,000, with a 0.7 m/z isolation width, 35% collision energy, 1 × 10^-5^ AGC target, and a maximum ion time of 86 ms. Dynamic exclusion was set to exclude previously sequenced peaks for 20 s within a 10-ppm (parts per million) isolation window. Only precursor ions with charge states between 2 and 6 were selected for fragmentation. For Set 2, elution was performed over a 30-minute gradient at a flow rate of 1250 nl/min, with buffer B/buffer (A+B) ratio ranging from 1% to 99% (buffer A: 0.1% FA in water; buffer B: 0.1% FA in 80% ACN). Mass spectrometry was conducted on a high-field asymmetric waveform ion mobility spectrometry (FAIMS) Pro-equipped Orbitrap Eclipse (Thermo) in positive ion mode. DDA employed 2 x 1.5-second top speed cycles and 2 compensation voltages (-45 and -65). Each top speed cycle comprised one full MS scan with as many MS/MS events as possible within the 1.5-second cycle time. Full MS scans were acquired at a resolution of 60,000 over an m/z range of 410 to 1600, with an AGC target of 4 × 10^-5^ and a maximum ion injection time of 50 ms. All HCD MS/MS spectra were collected at a resolution of 30,000, with TurboTMT on, a 0.7 m/z isolation width, 35% collision energy, 250% normalized AGC target, and a maximum ion time of 54 ms. Dynamic exclusion was configured to exclude previously sequenced peaks for 20 s within a 10-ppm isolation window. Precursor ions with charge states between 2 and 6 were selectively chosen for fragmentation.

### Database search parameters for TMT labeled samples

FP (version 18.0) was used to search both the discovery (Set 1) and replication (Set 2) datasets as essentially described ^41, 47^. First, mzML files were generated from the original MS .raw files (96 raw files/fractions per batch) of both Set 1 (3 TMT16 batches) and Set 2 (5 TMT18 batches) using the ProteoWizard MSConvert tool (version 3.0) with options including: ‘Write index’, ‘TPP compatibility’, and ‘Use zlib compression’, as well as “peakPicking” filter setting. Then all 96×8 mzML files from both sets were searched together using MSFragger (version 3.5). The human proteome database used comprised of 20,402 sequences (Swiss-Prot, downloaded 2/11/2019) and their corresponding decoys, including common contaminants. We also included the APOE2 and APOE4 coding variant: CLAVYQAGAR (APOE2), and LGADMEDVR (APOE4). Briefly, search settings included: Precursor mass tolerance was -20 to 20 ppm, the fragment mass tolerance was set to 20 ppm, mass calibration and parameter optimization were selected, and the isotope error was set to -1/0/1/2/3. The enzyme specificity was set to strict-trypsin and up to two missed cleavages allowed. Cleavage type was set to semi-enzymatic. Peptide length was allowed in the range from 7 to 35 and peptide mass from 200 to 5,000 Da. Variable modifications that were allowed in our search included: oxidation on methionine, N-terminal acetylation on protein, TMTpro modifications on serine, threonine and histidine as described ^48^, with a maximum of 3 variable modifications per peptide. Static modifications included: isobaric TMTpro (TMT16) modifications on lysine and the peptide N-termini as well as carbamidomethylationon of cysteine. MSFragger search results were processed using Percolator ^43^ for PSM validation, followed by Philosopher ^44^ for protein inference (using ProteinProphet ^49^) and FDR filtering. The reports of the quantified peptides and UniprotID-identified proteins with FDR < 1% were generated. All raw files, the database, the sample to TMT channel information, and the FP search parameter settings are provided on https://www.synapse.org/#!Synapse:syn52525880/files/.

All raw files from the consensus brain dataset ^11^, including 1080 raw files generated from 45 TMT 10-plexes for the Religious Orders Study and Memory and Aging Project (ROSMAP) BA9 tissues; 624 raw files generated from 26 TMT 11-plexes for ROSMAP BA6/BA37 tissues; 528 raw files generated from 22 TMT 11-plexes for the Banner Sun Health Research Institute (Banner) tissues; and 760 raw files generated from 20 TMT 11-plexes for Mount Sinai tissues, were re-searched using FragPipe (version 20.0) on the same Uniprot database as described above with slight modifications. This mainly included use of the first generation “TMT10” workflow rather than the newer generation TMTpro workflow and the cleavage type was set to enzymatic. Peptide length was allowed to range from 7 to 50. Variable modifications that were allowed in our search included: oxidation on methionine, N-terminal acetylation on protein, TMT10 modifications on serine, with a maximum of 3 variable modifications per peptide. Static modifications included: isobaric TMT10 modifications on lysine and peptide N-termini as well as carbamidomethylation on cysteine.

### Data normalization and variance correction for TMT labeled samples

The FP outputs of 8 batches from Set 1 and Set 2 or 113 batches from the consensus brain were integrated to get a combined raw abundance file. Protein levels were first scaled by dividing each protein intensity by the sum of all the reporter ion intensities of the TMT channel (each sample) followed by multiplying by the maximum channel-specific protein intensity sum. Proteins with more than 50% of missing values in samples were removed from the matrix prior to the further process (no imputation of missing values was performed). A tunable median polish approach (TAMPOR) was used to adjust technical batch variance as previously described ^50^. The algorithm is fully documented and available as an R function, which can be downloaded from https://github.com/edammer/TAMPOR. Following this, non-parametric bootstrap regression for batch within each cohort was performed for Set 1 and Set 2. For reprocessed brain dataset, we restricted our analysis to 456 samples of control, asymptomatic AD (AsymAD) and AD from the Banner (control = 26, AsymAD = 57, AD = 77, 22 TMT11 batches) and the ROSMAP (control = 75, AsymAD = 127, AD = 94, 36 TMT10 batches), and regressed for age, sex, post-mortem interval (PMI) and batch.

### Proteome coverage overlap and gene ontology (GO) enrichment analysis

All proteome overlap was visualized using the venneuler R package (v1.1-3) *venneuler* function. All functional enrichment was determined using the GOparallel function as documented on https://github.com/edammer/GOparallel. The Bader lab monthly updated .GMT formatted ontology gene lists ^51^ were used for retrieving GO annotation. Z-score and *p*-value from the one-tailed Fisher’s exact test (FET) followed by Benjamini-Hochberg (BH) FDR correction was used to assess the significance. A cutoff of Z-score > 1.96 (BH FDR corrected *p* < 0.05 and a minimum of five genes per ontology) was used as filter prior to pruning the ontologies.

### Protein differential abundance and hierarchical clustering

All differential abundance was presented as volcano plots that were generated with the ggplot2 package in R v.4.2.1. Pairwise differentially abundant proteins were identified using Student’s *t*-test, followed by Benjamini-Hochberg (BH) FDR correction. Supervised clustering analysis on differentially abundant Hp-enriched plasma proteins was performed with the R NMF package ^52^ in R v4.2.1. A cutoff of BH FDR-corrected *p* < 0.0005 was used to obtain 82 highly significant proteins and clustered with euclidian distance metric, complete linkage method using the *hclust* function is called from the NMF package *aheatmap* function.

### Correlation across platforms and replicate datasets

To evaluate the consistency across different platforms, we compared the Heparin-MS data with plasma measurements obtained using SomaScan® aptamer-based technology from SomaLogic (located in Colorado, USA) and proximity extension assay (PEA) technology from Olink® (based in Uppsala, Sweden). Data from SomaScan and PEA (referred to as “Olink” henceforth) were obtained for 35 (control = 18, AD = 17) out of the 36 individuals previously assessed using the Heparin-MS method. These data are accessible from a prior publication ^31^ and were subjected to cross-platform analysis using TAMPOR ^50^. Additionally, we performed correlation analyses between the two Heparin-MS sets (Set 1 and Set 2) using the log^2^ fold-change (AD vs Control) values. These correlation analyses were carried out and visualized employing the *verboseScatterplot* function from the R WGCNA package, utilizing the Pearson correlation coefficient and Student’s *p*-value to determine the statistical significance of these correlations.

### Meta-analysis of significance, Z-score transformation and correlation to AD biomarkers

Meta-analysis of significance on the combined selected samples (Set 1 + Set 2, *n* = 109) was performed using the R survcomp package *combine.test* function to calculate *meta p-*value. Average log^2^ fold-change (AD vs Control) between the two sets was used for the x-axis. Significantly altered proteins (*meta p-value* < 0.05) along with corresponding *meta p*-value and fold-change were listed in **Supplemental Table 16**. Variance-corrected protein abundance of all samples was then Z-transformed by subtracting the mean across 109 samples followed by dividing by standard deviation. Correlations and the Student’s *p*-value for their significance between Z-scores and immunoassay measures of AD-related traits, including cognition (MoCA score), CSF Aβ^1-42^, CSF tTau, CSF pTau181, CSF tTau/Aβ^1-42^ ratio and plasma pTau181, were calculated using the R WGCNA package *corAndPvalue* function.

### Protein re-assignment to consensus AD brain network modules

We reprocessed 456 of the Banner and ROSMAP brain samples used in our previously published WGCNA consensus brain network from 2022 ^11^, which underwent re-analysis with FP as outlined above. 8956 proteins were identified and the biweight midcorrelation (bicor) of each was calculated to the 44 eigenproteins of the original network, and a module assignment was made for the FP output proteins to the module with the highest positive correlation, if greater than or equal to 0.30, otherwise being assigned as “grey” ^53^. As described previously, a module eigenprotein represents the principal component of all proteins within a module. Moreover, we conducted bicor correlations and Student’s *p*-value for their significance to assess the association of module eigenproteins with Consortium to Establish a Registry for Alzheimer’s Disease (CERAD), Braak staging, and Mini-mental state examination (MMSE), using the *bicorAndPvalue* function from the R WGCNA package.

### Cell type enrichment analysis

As outlined previously ^21^, cell type enrichment for each module was performed by cross-referencing the corresponding gene symbols of each module with cell-type–specific gene lists derived from previously published RNA-seq data ^11, 54, 55^. Significance of cell type enrichment within each module was then determined using a one-tailed FET and corrected for multiple comparisons by the BH FDR method. The algorithm is fully documented and available as an R function, which can be downloaded from https://www.github.com/edammer/CellTypeFET.

### Overrepresentation analysis of differentially abundant plasma proteins in brain network

Hp-enriched plasma proteins that overlap with brain proteome and significantly altered in AD plasma compared to controls (Student’s *p* < 0.05) were assessed for overrepresentation in brain proteome using a one-tailed FET, and those modules with BH FDR-corrected *p* < 0.05 were considered significant. The background for this overrepresentation analysis comprised 8956 UniprotID-identified proteins from consensus brain ^11^ which was re-searched by FP as described above.

## Results

### Enrichment of heparin binding proteins from plasma

To assess the feasibility and efficacy of a heparin enrichment strategy from plasma, an equal volume (40 µl) of pooled human plasma, diluted into binding buffer, was introduced to heparin-sepharose resin in technical triplicate as described in our workflow (**Figure 1A**). Before performing MS analyses, fractions including diluted plasma (DP) inputs, the heparin-depleted flow-through (Hp-depleted FT), and the heparin-enriched (Hp-enriched) fraction, underwent gel electrophoresis and Coomassie Blue staining to visualize the proteins. These fractions were also prepared for immunoblotting to detect known HBPs, thrombin and APOE (**Figure 1B**). Immunoblotting demonstrated that heparin affinity chromatography was effective at enriching APOE and thrombin, which exhibited increased levels in the Hp-enriched fractions compared to the FT and DP inputs. Moreover, by Coomassie Blue staining we observed a reduction in the amount of albumin (∼66 kDa) in the Hp-enriched fractions compared to the inputs and FT. In support of these findings, single-shot label-free MS proteomic analyses revealed a notable disparity in the base peak chromatogram among the inputs, FT, and Hp-enriched fractions (**Figure 1C**). In the chromatogram spanning the 120-minute time window, the pattern of precursor peptide ion peak intensities in the input and FT fractions remained remarkably consistent, primarily dominated by a few prominent feature peaks, which is consistent with the presence of a highly abundant protein such as albumin. However, the base peak chromatogram for the Hp-enriched fractions exhibited much greater complexity, suggesting the presence of additional peptide species in the sample and a decrease in the dominance of the albumin peptides observed in the input and FT fractions. Consistently, following database search, we observed a significant reduction (∼2.5 fold) in the number of peptide spectral counts, a semi-quantitative measure of protein abundance, for albumin in the Hp-enriched fractions compared to the DP inputs and FT. Reciprocally, we observed a significant increase in the number of peptide spectral counts for thrombin (∼2.5 fold) and APOE (∼2.6 fold) in the Hp-enriched fractions compared to the DP inputs and FT, confirming our immunoblot findings (**Figure 1D**). Collectively, these observations suggest that the heparin-sepharose capture method not only enhanced the concentration of the HBPs, but also concurrently reduced the presence of highly abundant proteins like albumin in the Hp-enriched fraction. To evaluate the depth of protein identification across the fractions, we assessed the overlap of proteins that were detected in at least two out of three replicates of each fraction (**Figure 1E and Supplemental Table 1)**. Notably, the Hp-enriched fraction identified the greatest number of proteins (*n* = 771) compared to the DP input (*n* = 611) and FT (*n* = 527). Approximately 29% (221 out of the total 771 proteins) of the proteins in the Hp-enriched fraction were unique to this fraction. Gene ontology (GO) analysis conducted for the proteins identified in each fraction showed significant enrichment (Z-score > 1.3) of the term associated with ‘Heparin binding’ in the Hp-enriched fraction compared to those in the FT and DP inputs (**Figure 1F**).

**Figure 1.**
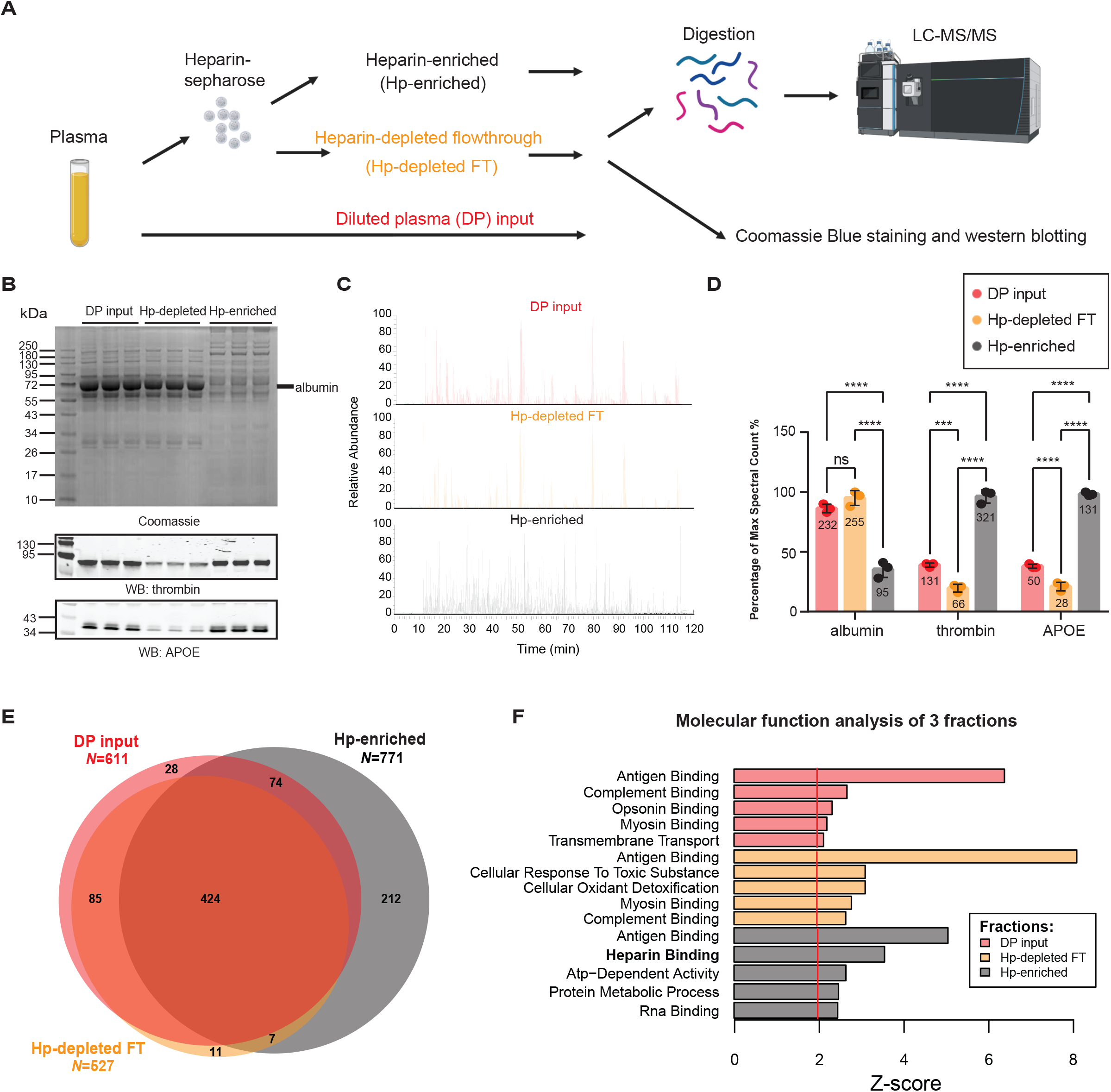
Heparin Enrichment of the plasma proteome. A) Step-by-step process of heparin enrichment and subsequent mass spectrometry (MS) analysis. Plasma samples were subjected to heparin enrichment, yielding three distinct fractions: DP input (*n* = 3), Hp-depleted FT (*n* = 3), and Hp-enriched fraction (*n* = 3). Each fraction underwent either Coomassie Blue staining, western blotting, or trypsin digestion before subsequent label-free MS analysis utilizing an Orbitrap Eclipse mass spectrometer. B) Coomassie Blue staining following western blotting was performed in triplicates for all fractions. A reduction in the amount of albumin (∼66 kDa) on the Hp-enriched fraction compared to the DP input and FT was illustrated by Coomassie Blue staining. Enrichment was also determined by western blotting for thrombin and APOE on the Hp-enriched fraction. C) MS base peak chromatograms of each fraction reveal the variation in sample complexity. The Hp-enriched fraction exhibits a notable increase in sample complexity compared to the DP input and Hp-depleted FT fractions, reflecting successful enrichment of low abundance proteins in the Hp-enriched fraction. D) Total number of peptide spectral counts (reported as the maximum percentage) for albumin, thrombin, and APOE in each fraction. Actual spectral counts are labeled within columns. Albumin is significantly depleted in the Hp-enriched fraction compared to the DP input. Conversely, thrombin and APOE are significantly enriched in this fraction. ANOVA with Tukey post-hoc correction was used to determine the *p*-values (*\* p* < 0.05, ** *p* < 0.01, *** *p* < 0.001, **** *p* < 0.0001). E) The number of proteins identified in each fraction: DP input (*N* = 611, Hp-depleted FT (*N* = 527), and Hp-enriched fraction (*N* = 771) and the degree of protein overlaps among them. F) GO terms for proteins detected in each fraction, using 841 total proteins measured in at least 2 out 3 replicates within each fraction as background, highlighting the enrichment of proteins associated with the “Heparin binding” molecular function in the Hp-enriched fraction. Z-score > 1.3 (*p* < 0.05) is significant. WB, western blotting; LC-MS, liquid chromatography-mass spectrometry.

To comprehensively evaluate the extent of heparin enrichment, we utilized the protein abundance values based on MS precursor signal intensities from both the DP inputs (*n* = 3), Hp-depleted FT (*n* = 3) and the Hp-enriched fractions (*n* = 3) where we applied a significance threshold of *p* < 0.05 and a fold-change threshold of > 2 across 821 proteins. In total, 518 proteins exhibited significant changes in their abundance levels across the two fractions (**Supplemental figure 1A and Supplemental Table 2**), with 338 increased and 180 decreased, when comparing the Hp-enriched fractions to the DP inputs. Among these, we consistently observed the enrichment of key HBPs such as thrombin (F2), APOE, and APP, which possesses a heparin binding domain^56^. Additionally, other neurodegeneration related proteins such as neurogranin (NRGN) and valosin-containing protein (VCP), also exhibited significant enrichment in the Hp-enriched fractions. As expected, albumin (ALB) was significantly decreased in the Hp-enriched fractions in addition to other highly abundant proteins decreased including transferrin (TF), A2M and ORM, SERPINA1, and APOA1. Notably, APOE has three genetic variants (*ε* 2, *ε* 3, *ε* 4) associated with AD risk: APOE4 has the highest risk, APOE2 the lowest risk, and APOE3 the intermediate risk ^22^. APOE4 and APOE2 variants can be identified through coding changes that result in isoform specific peptides following trypsin digestion^57^. In the pooled plasma sample used here, all APOE protein variants were detected and quantified. Interestingly, APOE4 showed the most enrichment in Hp-enriched fractions (15.5-fold, *p* = 0.0003), followed by APOE (14.7-fold, *p* = 0.0004), which was inferred as APOE3 due to a lack of variant specific peptides, and APOE2 (7.4-fold, *p* = 0.006) (**Supplemental Figure 1B**). This aligns with prior research, highlighting differences in APOE heparin affinity between the variant isoforms (APOE4 > APOE3 > APOE2) ^58, 59^, suggesting that the genetic susceptibility to AD attributed to APOE is, in part, linked to heparin binding. Collectively, our findings support successful enrichment of not only APOE, but other HBPs from human plasma.

### Heparin-enriched plasma proteome is significantly altered in AD

After successfully demonstrating enrichment of HBPs from human plasma, we aimed to both enhance the depth and determine the differences in the plasma proteome between AD (*n* = 18) and control (*n* = 18) individuals. Following heparin affinity enrichment, we employed tandem mass tag mass spectrometry (TMT-MS) in conjunction with high-pH off-line fractionation to identify a total of 3284 proteins (**Figure 2A and Supplemental Table 3**). AD diagnoses were established based on notable cognitive impairment as determined by the Montreal Cognitive Assessment (MoCA), with scores averaging 13.1± 7.2 for AD and 27 ± 1.6 for controls (**Table 1**). These diagnoses were further supported by the presence of low Aβ levels and elevated tTau and pTau levels detected in the CSF immunoassays for AD and normal levels of these biomarkers in controls (**Table 1**). Furthermore, we measured pTau181 in the plasma samples which showed significant differences between control and AD cases (**Figure 2B**). Prior to subjecting the samples to TMT-MS, we confirmed the heparin enrichment of APOE and thrombin via immunoblotting of the input, FT, and Hp-enriched fractions (**Supplemental Figure 2**). To enhance data completeness, we considered only those proteins quantified in at least 50% of the samples for subsequent analyses, culminating in the final quantification of 2077 proteins (**Supplemental Table 5**). To identify differentially abundant proteins in the AD plasma proteome, we generated a volcano plot for pairwise comparisons of AD versus control individuals (**Figure 2C**). Proteins with significant changes in abundance within the AD group were determined using a Student’s *t*-test (*p* < 0.05). The complete list of differentially abundant proteins is provided in **Supplementary Table 6**. We identified 579 proteins with significantly increased abundance and 661 proteins with significantly decreased abundance in AD cases. Notably, our findings support an increase in M42 matrisome-associated proteins in both AD brain and plasma, such as SMOC1 (*p* = 2.26 × 10^-6^), OLFML3 (*p* = 1.06 × 10^-6^), APOE (*p* = 0.028611), MDK (*p* = 0.000367), SPON1 (*p* = 3.07 × 10^-5^), GPNMB (*p* = 0.000935), and FRZB (*p* = 0.000555) ^11, 31^. Particularly noteworthy is the identification of SMOC1, MDK, GPNMB, and FRZB, which had not previously been measured in plasma using MS-based technology ^31^. Additionally, we discovered other highly significant proteins, including SMOC2 (*p* = 4.15 × 10^-6^), APOE4 (*p* = 0.000698), BGN (*p* = 1.68 × 10^-7^), ESM1 (*p* = 1.40 × 10^-7^), CSF1 (*p* = 7.52 × 10^-6^), PLA2G7 (*p* = 0.004595), GCG (*p* = 0.004995), and PODN (*p* = 2.25 × 10^-8^). As expected, levels of APOE4 were more significant in the AD group compared to the control group, consistent with the higher frequency of APOE4 carriers in AD. GO analysis of the 579 significantly increased proteins in AD indicated that ‘Heparin binding’ was a major altered pathway (**Figure 2D**), which aligns with the strong correlation observed between M42 members and HBPs in AD brain ^11^. Conversely, the 661 significantly decreased proteins were strongly linked to ‘ATP binding’ and ‘Mitotic cell cycle’ processes among others. To investigate differential expression at the individual sample level, we used proteins with BH FDR-corrected *p* < 0.0005 to perform supervised cluster analysis across all 36 samples. As illustrated in **Figure 2E**, the expression profiles of 82 highly significant proteins in Hp-enriched plasma correctly distinguished AD from control cases, with only minor exceptions. Among these 82 proteins, 48 were increased in AD, while 34 were decreased. In summary, these results revealed hundreds of differentially abundant plasma HBPs that are altered in AD.

**Figure 2.**
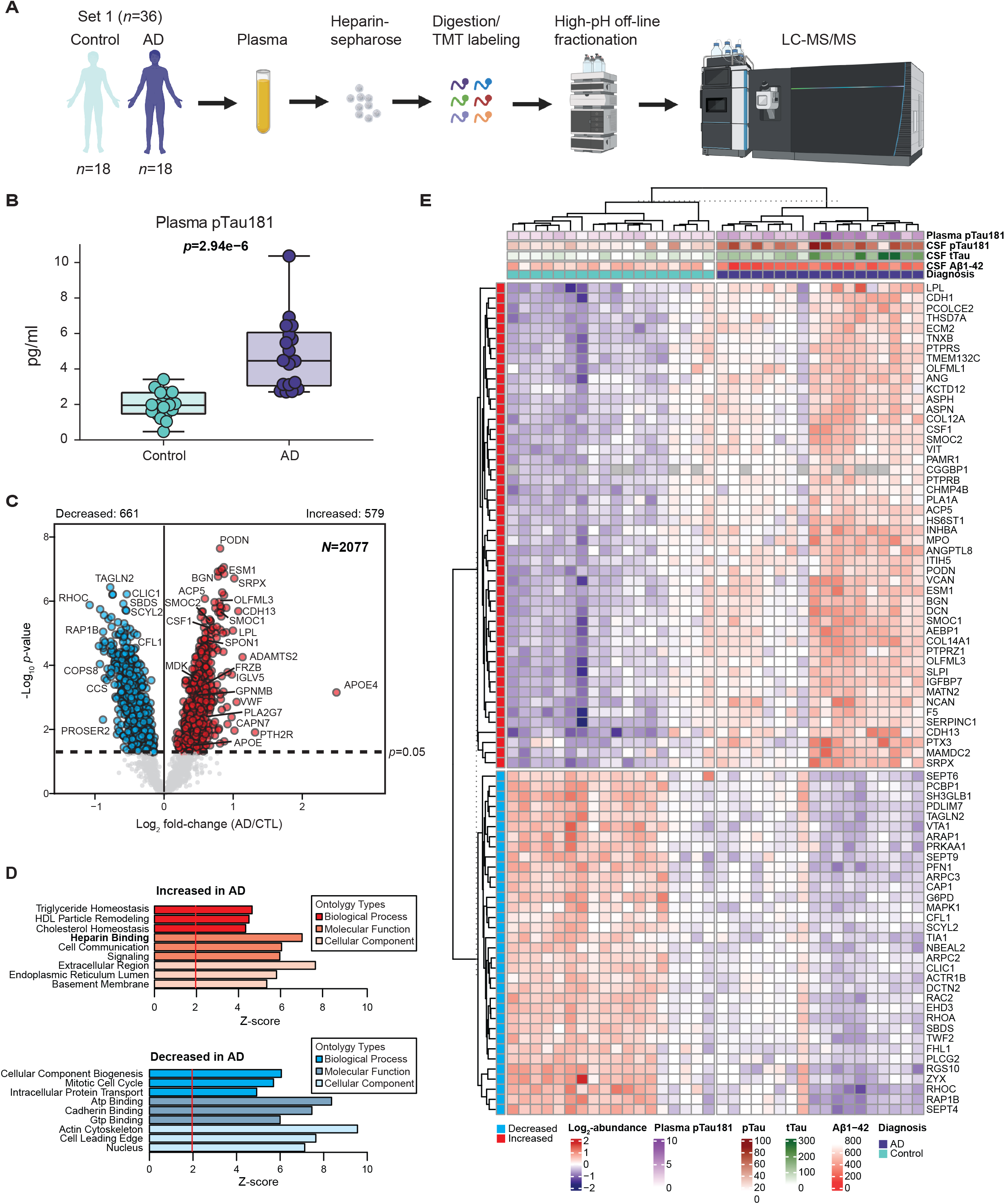
Heparin-enriched plasma proteome is significantly altered in AD. A) Plasma samples were collected from both the control group (*n* = 18) and individuals with AD (*n* = 18). These samples underwent heparin-sepharose enrichment, followed by trypsin digestion and TMT labeling. Subsequently, high-pH off-line fractionation and LC-MS/MS analysis were conducted using an Orbitrap Lumos mass spectrometer. B) Measurements of pTau181 across all 36 samples were displayed. Statistical significance was determined by Student’s *t*-test (*p* = 2.94 × 10^-6^). C) The volcano plot illustrates the differential abundance of 2,077 proteins between the control and AD groups. The x-axis represents the log^2^ fold-change (AD vs CTL), while the y-axis represents the Student’s *t*-statistic (-log^10^ *p*-value) calculated for all proteins in each pairwise group. Proteins significantly increased in AD (*N* = 579) are highlighted in red (*p* < 0.05), whereas those significantly decreased in AD (*N* = 661) are depicted in blue. Grey dots represent proteins with unchanged levels. D) Top GO terms in the 579 increased (red) or 661 decreased (blue) proteins in AD measured in Set 1 considering the background of 2077 proteins in the proteome. Three GO terms with the highest Z-scores within the domains of biological process, molecular function, and cellular components are presented. E) A supervised cluster analysis was conducted across the control and AD plasma of discovery samples (Set 1), employing the 82 most significantly altered proteins in the dataset (BH FDR-corrected *p* < 0.0005). pTau, phospho-tau; tTau, total tau; CTL, control; LC-MS, liquid chromatography-mass spectrometry.

### Heparin-enriched AD plasma proteome demonstrates consistency and complementarity to other independent proteomic platforms

To assess both the validity and the consistency of the direction of change in our Hp-enriched AD plasma proteome, we utilized protein measurements, previously obtained through either the SomaScan® aptamer-based technology or the proximity extension assay (PEA) technology from Olink®, from overlapping cases in our discovery set (Set 1) of control and AD plasma samples (**Figure 3**). The protein measures were obtained from these two platforms following cross-platform TAMPOR as previously described ^31^ and are provided in **Supplemental Table 7 and 8**. As anticipated, the aptamer-based SomaScan yielded the largest set of protein measurements from the plasma samples (N = 7284), followed by our Heparin-MS method (N = 2077) and Olink (N = 979). We used gene symbols from each output to calculate the number of measurements and their overlap across the various platforms (**Figure 3A and Supplemental Table 9**). Each platform exhibited its own unique group of non-overlapping identified proteins (gene products), which we subsequently subjected to GO analyses to assess their associations with biological pathways. Notably, the unique gene products identified in the Heparin-MS method exhibited significant association with pathways related to ‘Alzheimer’s disease and miRNA effects’ and ‘Parkin-ubiquitin proteasomal system pathway’ **(Figure 3B**), which further underscores the value of heparin enrichment in capturing biology related to AD and related neurodegenerative diseases. To gauge the consistency in the direction of change in AD plasma proteome compared to controls across independent platforms, we conducted correlation analyses of the log^2^ fold-change (AD vs Control) for overlapping proteins between Heparin-MS and Olink (*N* = 279 proteins, *cor* = 0.73, *p* = 1.1e^-47^) or SomaScan (*N* = 1183 proteins, *cor* = 0.62, *p* = 1.4e^-126^). Both comparisons revealed strong positive correlations between the two platforms (**Figure 3C**). It is worth noting that no enrichment method was employed in the Olink or aptamer-based measurements. However, the aptamer-based and in some instances the Olink assays necessitate multiple separate dilutions to accommodate the wide dynamic range of plasma protein concentrations. Thus, these findings indicate that the observed direction of change in Hp-enriched plasma by TMT-MS in the AD group versus the control group largely aligns with the levels measured in the unprocessed plasma inputs. Notably, increases of several M42 members in AD were consistently validated by at least two platforms, including SMOC1, OLFML3, GPNMB, HTRA1, and APOE, whereas SPON1, PTN, APP and FRZB showed the same direction of change in AD across all three platforms. Additional Hp-enriched plasma proteins such as ESM1, PLA2G7, BGN, CSF1, GCG, VWF, and LPL were also consistently increased in AD across at least two platforms. Boxplots for SPON1, WAS, ESM1 and PLA2G7 that showed concordant changes in all three platforms are provided (**Figure 3D**). This data substantiates a robust overall correlation among the three proteomic platforms regarding the directional changes observed in AD plasma. Furthermore, it suggests that heparin enrichment of the plasma proteome does not significantly bias the direction of change in AD. Overall, these data support the validity and consistency of the Hp-enriched plasma proteome in AD across independent platforms.

**Figure 3.**
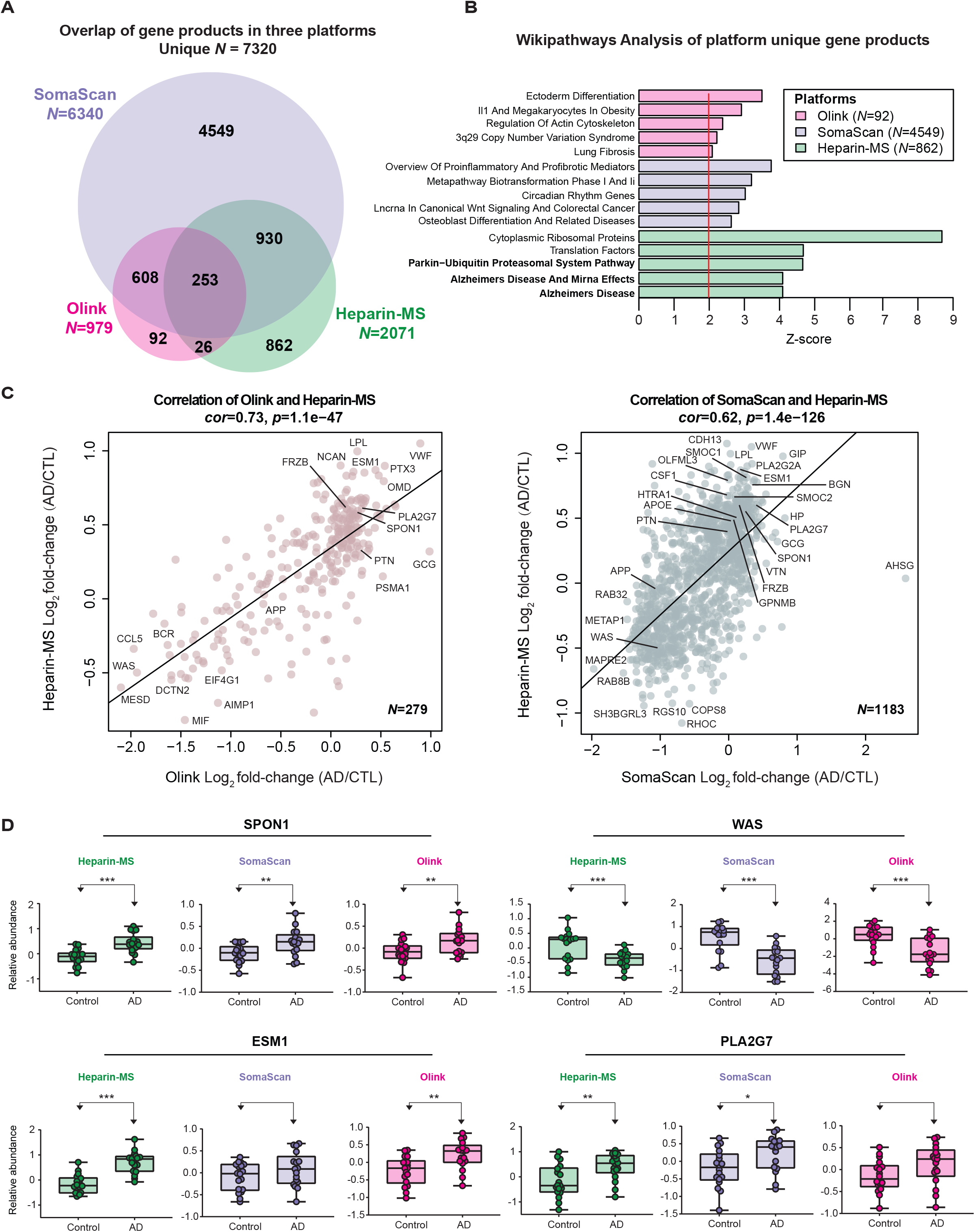
Heparin-enriched plasma proteome is consistent and complementary to other independent proteomic platforms. A) The number and overlap of proteins (i.e., unique gene products) quantified in plasma across three platforms, including the TMT-MS approach in Hp-enriched samples (Heparin-MS, control = 18, AD = 18), the PEA-based assay (Olink, control = 18, AD = 17), and the aptamer-based method (SomaScan, control = 18, AD = 17). The inclusion criterion required at least 18 measurements across all samples. B) Wiki-pathway analysis highlighting specific pathways from uniquely identified gene products in each of the three platforms. The Heparin-MS method exhibited significant association with pathways related to ‘Alzheimer’s disease and miRNA effects’ and ‘Parkin ubiquitin proteasomal system pathway’, underscoring the neurodegenerative disease specificity. A total of 5503 platform-unique gene symbols were used as background for GO analysis. C) Pearson correlation between log^2^ fold-change (AD vs CTL) of common gene products measured by the Heparin-MS method and Olink (left, *N* = 279 gene products, *cor* = 0.73, *p* = 1.1e^-47^) as well as the Heparin-MS method and the SomaScan (right, *N* = 1183 gene products, *cor* = 0.62, *p* = 1.4e^-126^). The significance of Pearson correlation was determined by Student’s *p*-value. Several M42 members and associated proteins showed concordant directions in both comparisons, including SPON1, APP, PTN, FRZB, ESM1, PLA2G7, VWF, GCG, and LPL. D) Boxplots display the consistent and statistically significant changes in AD observed for SPON1, WAS, ESM1, and PLA2G7 across all three platforms. Significance was determined by Student’s *t*-test (*\* p* < 0.05, ** *p* < 0.01, *** *p* < 0.001). CTL, control; *cor*, Pearson correlation coefficient.

### Heparin enrichment enhances the depth of plasma proteome and is reproducible for abundance changes in AD

To demonstrate the consistency and reproducibility of the plasma heparin enrichment approach, we analyzed a separate set of samples (Set 2) involving 49 controls and 36 AD individuals (**Supplemental Table 10**), which included 13 overlapping controls with the discovery Set 1. These samples underwent heparin enrichment processing similar to the procedures applied to Set 1, albeit using a different volume and lot of heparin-sepharose beads (**Figure 4A**). TMT-MS was employed for the analysis of the Hp-enriched fraction, carried out across five batches, resulting in the identification of 2618 proteins measured in 50% or more of the samples. (**Supplemental Table 11**). Collectively, the two datasets identified a total of 3284 unique proteins but only 2866 in 50% or more of the samples within each set were used for further analysis (**Figure 4B**). This number is more than 2-fold greater than previous coverage generated with immune-depletion method ^31^, supporting the depth and sensitivity of the heparin enrichment approach when coupled to TMT-MS. Impressively, this encompassed proteins across 10 orders of magnitude in concentration within the plasma, even extending to those with the lowest concentration (1.47 pg/ml), as evidenced by inclusion of proteins such as LAG3 and RNF213 as well as many M42 matrisome members (**Figure 4C and Supplemental Table 12**). In total, 1829 out of 2866 proteins overlapped between the two independent TMT-MS datasets (**Figure 4D and Supplemental Table 13**). Differential expression analyses were performed on all 2618 proteins within Set 2, resulting in 826 proteins being increased and 795 proteins decreased in AD (**Supplemental Table 14**). Notably, significant protein changes between AD and control samples across the two sets were highly reproducible, (**Figure 4E)**. Among the 732 overlapping proteins with a BH FDR-corrected *p*-value < 0.05 in both datasets, only 10 (1.4%) exhibited discordant changes in the two sets (*cor* = 0.93, *p* < 1e^-200^). However, when increasing the significant threshold to *p* < 0.01, we observed no discordant difference between the 390 overlapping proteins. Among these highly significant proteins, we confirmed several targets outside of M42 matrisome members that were consistently validated across different proteomic platforms (**Figure 3**), including ESM1, PLA2G7, BGN, CSF1, and GCG, further reinforcing their changes in AD plasma. Overall, our heparin enrichment approach coupled to TMT-MS consistently and reliably captured a diverse spectrum of plasma proteins, spanning an impressive dynamic range of magnitude.

**Figure 4.**
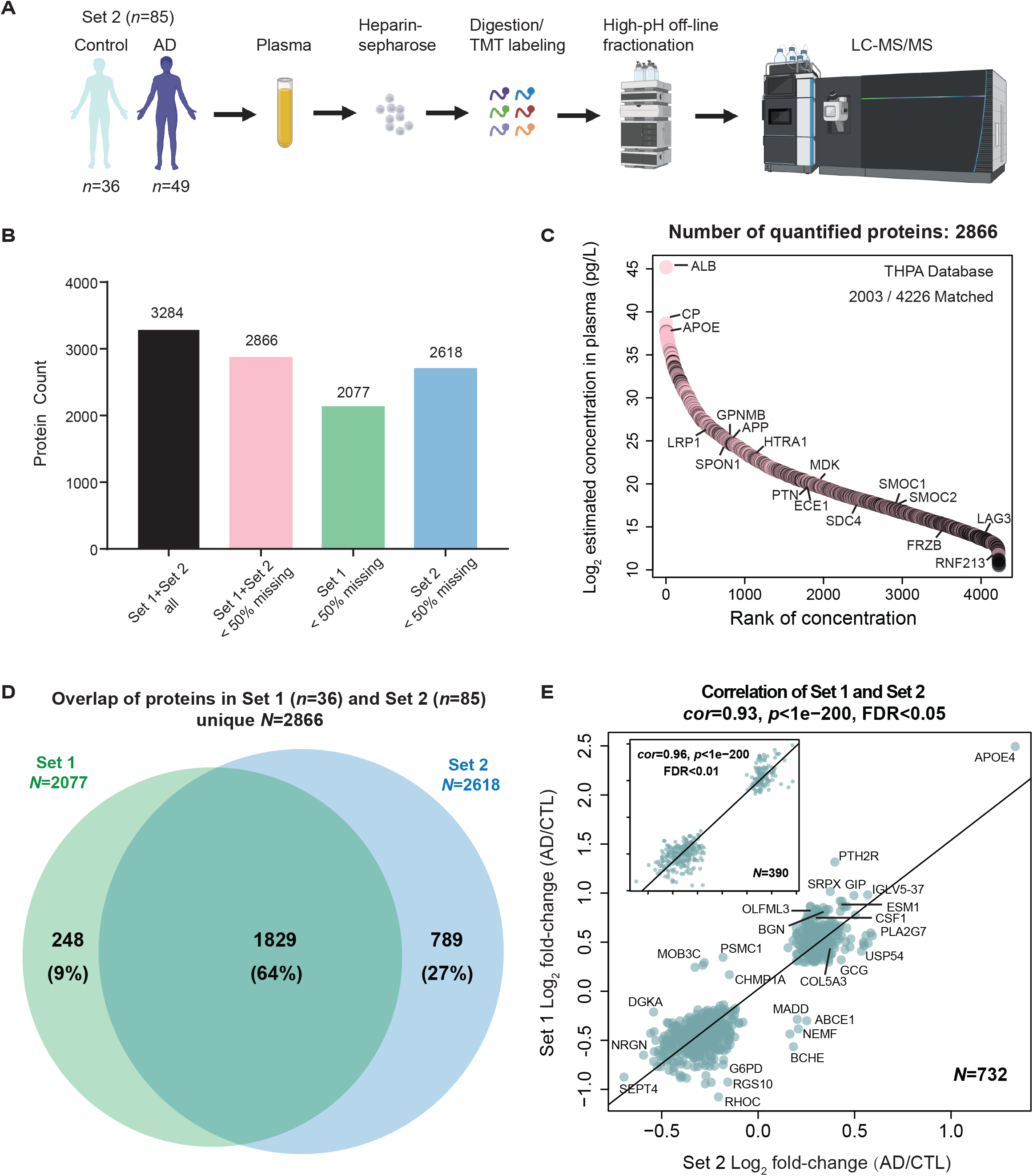
Heparin enrichment enhances the depth of plasma proteome and is reproducible in AD. A) The second replication dataset (Set 2) was comprised of control (*n* = 36) and AD individuals (*n* = 49), which underwent similar heparin enrichment and TMT-MS analysis as the discovery dataset (Set 1). B) The combined datasets yielded a total of 3284 unique proteins, and 2866 of them being identified and quantified in 50% or more of the samples (< 50% missing) within each set. Set 1 and Set 2 identified 2077 and 2618 proteins respectively, with 50% missing. C)The 2866 proteins were ranked by their log^2^ estimated concentration (pg/L) in plasma. Protein concentration information was obtained from The Human Protein Atlas Database (see methods). Notably, this set of proteins covered an impressive range of concentrations, spanning ten orders of magnitude. Even proteins with the lowest concentration, such as LAG3 and RNF213, were included, along with numerous members of the M42 matrisome that are highlighted. D) The number and overlap of proteins quantified in Set 1 (*N* = 2077) and Set 2 (*N* = 2618) with less than 50% missing values. Among the 2866 total unique proteins identified, approximately 64% (1829 out of 2866) were overlapping between the datasets. E) A scatter plot illustrates the Pearson correlation between log^2^ fold-change (AD vs CTL) of significantly altered proteins in both Set 1 and Set 2. There’re 732 proteins overlapping in the two sets and being significantly changed in AD with a BH FDR-corrected *p*-value < 0.05 in both datasets. Only 10 out of 732 proteins exhibited discordant changes in the two sets, demonstrating a high degree of concordance (*cor* = 0.93, *p* < 1e^-200^). Furthermore, all 390 proteins selected with a BH FDR-corrected *p*-value < 0.01 displayed consistent changes in both datasets when comparing AD and control samples, with a remarkable correlation of 0.96 (*p* < 1e^-200^). The significance of Pearson correlation was determined by Student’s *t*-test. CTL, control; LC-MS, liquid chromatography-mass spectrometry; *cor*, Pearson correlation coefficient.

### Association of heparin binding proteins in plasma to cognitive measures and conventional AD biomarkers: CSF Aβ^1-42^, tTau and pTau181 and plasma pTau181

CSF biomarkers, including Aβ^1-42^, tTau, and pTau181, have played a pivotal role in identifying individuals who are either at risk or already manifesting underlying AD pathology, collectively referred to as AT+ individuals ^35^. Furthermore, recent advancements have brought plasma pTau species, specifically pTau181, pTau231 and pTau217, into focus due to their associations to both underlying amyloid and tau pathology, even during the preclinical stages of the disease ^60, 61^. Here, we aimed to evaluate proteins within the two sets of Hp-enriched plasma proteome that are associated with cognition and AT+ status by comparing their abundances to MoCA scores, CSF AD biomarkers (Aβ^1-42^, tTau, tTau/Aβ^1-42^, pTau181) and plasma pTau181 collected on the same patients. To enhance the specificity of our analysis, we excluded cases from the initial 121 samples (including 13 overlapping controls as described above) that did not meet strict CSF biomarker criteria (tTau/Aβ^1-42^ ratio) or MoCA cutoffs ^35, 36^ as outlined in **Supplemental Figure 3A**. As a result of this selection process, 12 non-overlapping samples were excluded from Set 2, resulting in a total of 109 samples (Set 1 = 36 samples, Set 2 = 73 samples) for further analysis (**Supplemental Table 15)**. Boxplots for each measurement across all unique cases were generated to show their separation based on disease status (**Supplemental Figure 3B**). Subsequently, we employed a meta-analysis of Set 1 and Set 2 to generate a composite *p*-value that considered the significance of association and effect size (AD vs Control) for 2865 proteins, all of which exhibited less than 50% missing values in either dataset. This meta-analysis revealed a substantial increase in differentially abundant proteins in the Hp-enriched AD plasma, with 945 proteins showing an increase and 923 proteins showing a decrease in AD, as compared to analyzing either dataset alone. **(Figure 5A, Supplemental Table 16)**. Importantly, this approach also ensured the retention of proteins that were identified in only one dataset (e.g., SMOC1 in Set 1), thereby preserving their availability for subsequent analyses. To assess the relationship between Hp-enriched plasma proteins to AD biomarkers and cognitive measures, we conducted correlation analyses between Z-transformed protein abundance (**Supplemental Table 17**) and immunoassay values of CSF (Aβ^1-42^, tTau, pTau181, tTau/Aβ^1-42^), plasma pTau181, and MoCA scores (**Supplemental Table 18)**. In **Figure 5B**, we highlight 77 Hp-enriched plasma proteins that exhibit significant correlations with at least three of the measurements (AD biomarkers or MoCA scores). Individual correlation scatterplots and linear fit lines are presented for ESM1, BGN, PLA2G7, and CSF1, all of which exhibit significant correlations with AD biomarkers and cognitive measures (**Figure 5C**). Moreover, these proteins have consistently demonstrated increased abundance in AD plasma in both Set 1 and Set 2 as well as other platforms as described above. Therefore, the correlation between these Hp-enriched plasma proteins to established AD biomarkers and cognitive measures suggests the potential utility of these proteins to be used for disease classification or staging in the context of AD progression.

**Figure 5.**
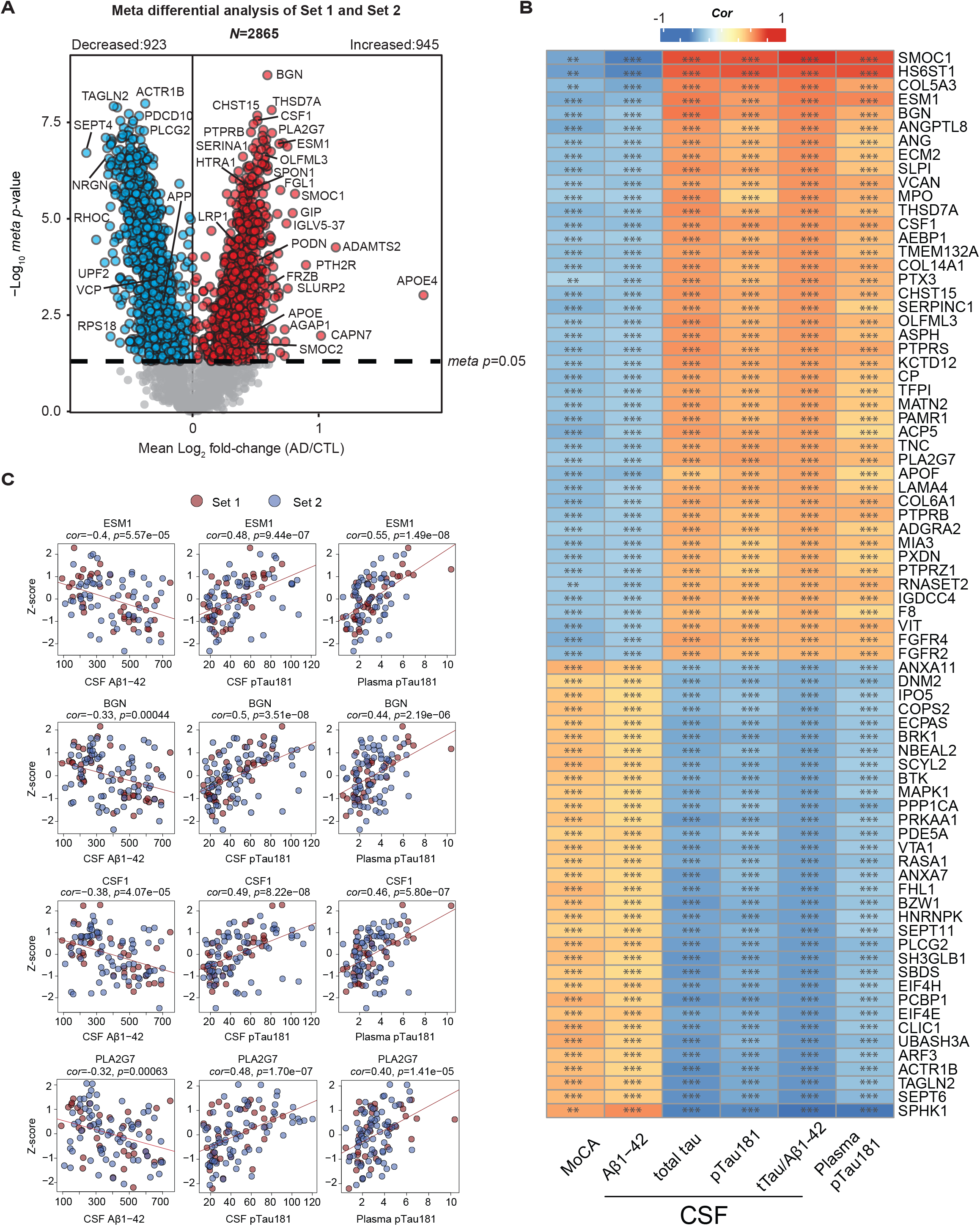
Association of heparin binding proteins in plasma to conventional AD biomarkers: CSF Aβ^1-42^, tTau and pTau181 and plasma pTau181. A) A meta-analysis of significant differences between control and AD on 2865 Hp-enriched plasma proteins that were measured in 50% or more samples within filtered Set 1 (control = 18, AD = 18) and Set 2 (control = 29, AD = 44). The x-axis represents the mean log^2^ fold-change (AD vs CTL), indicating an average abundance difference between Set 1 and Set 2. The y-axis shows the Student’s *t*-statistic (-log^10^ *meta p*-value) for all proteins in each pairwise group. Significantly increased proteins in AD are marked in red (*p* < 0.05), while proteins with significantly decreased levels in AD are denoted in blue. Grey dots represent proteins with unchanged levels. B) The heatmap highlights 77 Hp-enriched plasma proteins that have strong correlations to AD biomarkers. The color scale represents the degree of Pearson correlation (positive in red and negative in blue) between Z-transformed plasma protein abundances and immunoassay measures of various AD-related traits, including cognition (MoCA score), CSF Aβ^1-42^, CSF tTau, CSF pTau181, CSF ratio of tTau/Aβ^1-42^, and plasma pTau181. Significance levels determined by Student’s *t*-test are denoted by overlain asterisks; \**p* < 0.05, \*\**p* < 0.01, \*\*\**p* < 0.001. C) Individual scatterplots illustrate the correlations with CSF Aβ^1-42^, CSF pTau181, and plasma pTau181 of four specific Hp-enriched plasma proteins: ESM1, BGN, CSF1, and PLA2G7. *Cor* and *p*-values for each correlation are provided above each plot. The colors are differentiated by sets, with red representing Set 1 and blue denoting Set 2. CTL, control; *cor*, Pearson correlation coefficient.

### Overlap between the heparin-enriched plasma and human brain proteome

Previously integrated analysis of the human brain and CSF proteomes has revealed a substantial overlap of approximately 70%, strongly supporting a hypothesis that CSF serves as a valuable window into the brain ^14^. However, less is known between the overlap in plasma and brain proteomes. While studies have explored the overlap between brain and plasma proteomes using TMT-MS proteomic datasets following the immune-depletion of highly abundant proteins ^31^, we sought to leverage the depth of our Hp-enriched TMT-MS proteomic datasets to assess the overlap with human postmortem brain proteome. To ensure the consistency of protein overlap analysis, we employed the same Uniprot database and utilized the FragPipe search algorithm to re-analyze 456 dorsolateral prefrontal cortex (DLPFC) tissues from control, asymptomatic AD (AsymAD) and AD brains from the ROSMAP and the Banner cohorts (**Supplemental Figure 4A and Supplemental Table 19**) ^11^. The classification of cases was established according to a standardized classification system, which incorporated semi-quantitative histopathological assessments of Aβ and tau neurofibrillary tangle deposition, along with an evaluation of cognitive function in proximity to the time of death, as detailed in our previous work ^11^. Specifically, AsymAD cases were characterized by a neuropathological burden of Aβ plaques and tau tangles that closely resembled AD cases. However, these individuals did not exhibit substantial cognitive impairment near the time of death, indicating an early preclinical stage of AD. By applying a data normalization approach similar to the one used for the Hp-enriched plasma samples, we successfully identified 8,956 proteins detected in 50% or more of the samples from brain tissues corresponding to 8,904 unique gene symbols (**Supplemental Table 20**).When compared to a previous brain proteome dataset generated using Proteome Discoverer (PD) and an older 2015 Uniprot protein database ^11^, it was found that 7,476 of these unique gene symbols overlapped between the two outputs, while 1,428 were exclusive to the FragPipe, representing an approximately 18% increase in the number of identifications (**Supplemental Figure 4B and Supplemental Table 21**). This included proteins such as SMOC2, which is highly homologous to SMOC1 ^62^ and also increased in AD brain and plasma (**Figure 2C**). Furthermore, to ensure the consistency in the effect size of the AD brain proteome compared to controls, we correlated the differentially abundant proteins (**Supplemental Table 22 and Supplemental Table 23**) across these two sets of results, revealing a correlation coefficient of 0.9 with a *p*-value <1e^-200^ between AD and Control groups and a correlation coefficient of 0.84 with a *p*-value < 1e^-200^ between AsymAD and Control groups (**Supplemental Figure 4C**). This robust correlation provides strong support for the reliability of the FragPipe quantitative output. Subsequently, we systematically mapped all the measured brain proteins by FragPipe to one of the 44 pre-existing network eigenproteins from a consensus brain network ^11^. This mapping was achieved by recalculating the kME (bicor correlation to module eigenprotein) for each protein and assigning it to the module with the highest correlation (**Supplemental Table 24**). A revised M42 was generated (**Figure 6A**), comprising 35 members, marking a 10% increase compared to the number observed in the previous TMT-MS brain dataset using PD ^11^. This new M42 included additions such as SMOC2, COL5A1, SLIT3, HGF, NXPH2, RSPO2, and CHADL, among which SMOC2 and COL5A1 exhibited significant increases in AD plasma levels. Moreover, the incorporation of APOE-specific protein isoforms into the database led to the assignment of APOE4 to M42, rather than APOE (inferred as APOE3), following the FragPipe search of the brain proteome. Although APOE exhibited the strongest association with M42 (kME = 0.287) compared to all other modules in the network, it did not meet the 0.3 kME cutoff for inclusion into a network module. This finding further bolsters the genetic connection between the *APOE* ε4 genotype and increased levels of M42 members ^11,63^. Another four previous members of M42 that showed no change in plasma levels were reassigned to other modules. These members included ECE1, COL11A1, QPRT, and FLT1, which exhibited correlation with M42, yet were assigned to other modules within the network, likely due to their centrality as bottleneck nodes (**Supplemental Table 24**). When comparing the proteins identified in plasma with those in the brain proteome, we discovered that a substantial portion, approximately 77% (2211 out of 2685), of the plasma-identified proteins were also quantified in the brain (**Figure 6B and Supplemental Table 25**). Surprisingly, this was a higher percentage than we have reported for the CSF-brain overlaps (∼70%) ^14^. However, it is important to note that many of these proteins in the Hp-enriched plasma are likely not exclusively derived from the brain, as they are ubiquitously expressed throughout the body. To assess the overlap between the brain network and the plasma proteome, we mapped Hp-enriched plasma proteins to one of the 44 brain co-expression modules in **Figure 6B**. Among these modules, those associated with M40 ‘Antigen binding’ (with a 98% overlap of members) and M26 ‘Acute phase response’ (with an 89% overlap of members) exhibited the highest degree of overlap with the plasma proteome. Notably, M42 ‘Matrisome’, which exhibited the most robust correlation with AD pathology in the brain ^11, 31^, demonstrated a 46% overlap (16 out of 35 members) with the plasma proteome. This represents a significant improvement compared to the immune-depletion approach, which resulted in the overlapping of only 8 out of 32 members ^31^, reflecting a mere 25% of the module.

**Figure 6.**
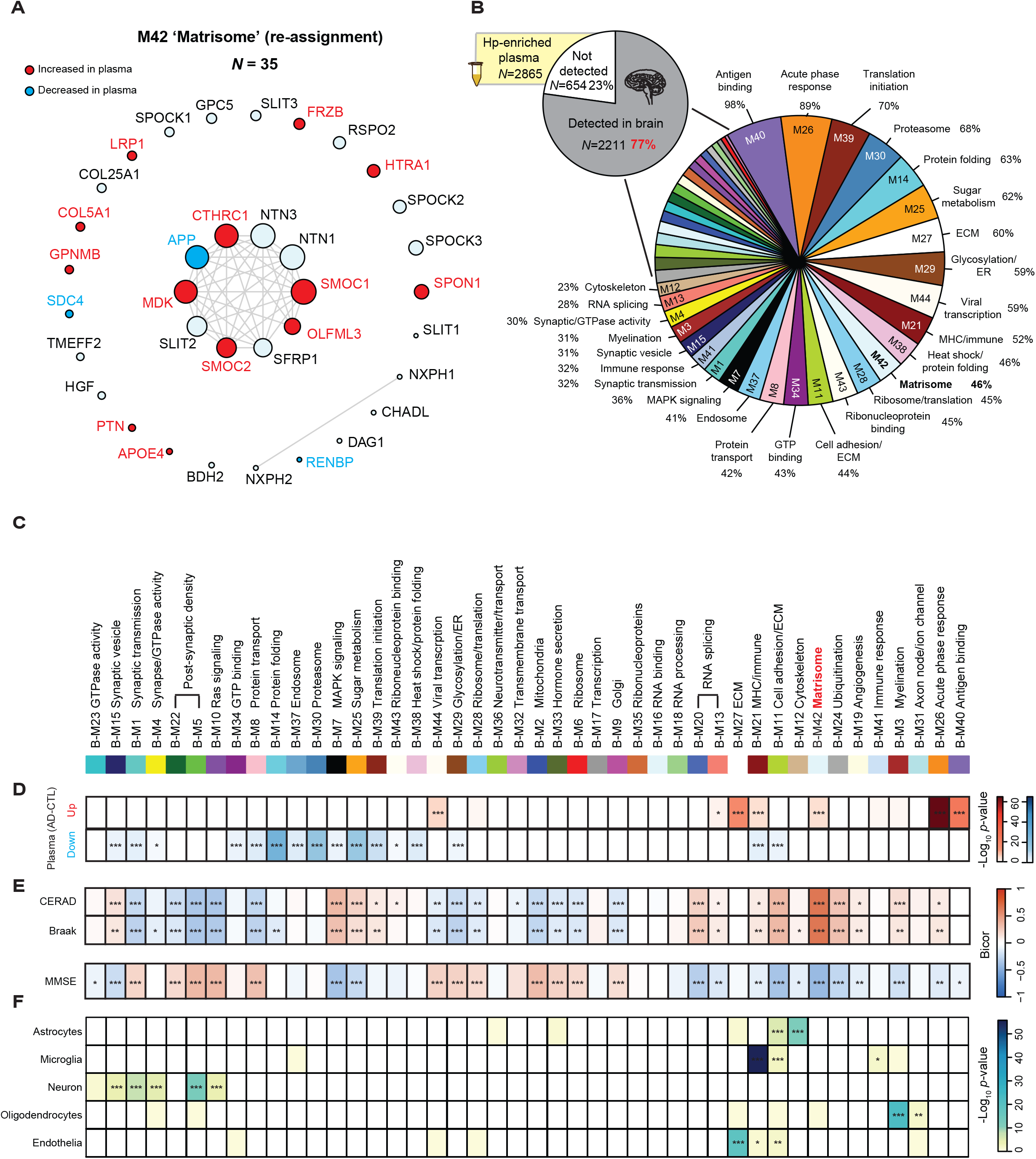
Mapping differential abundant heparin-enriched plasma proteins in AD within a human consensus brain network. The I-graph displays the updated M42 membership following FP database search, which includes 35 total proteins. Members with increased abundance in Hp-enriched AD plasma are highlighted in red, while those with decreased abundance are indicated in blue. B) The pie chart shows the number of proteins that overlap between the Hp-enriched plasma dataset (*N* = 2865) and consensus human brain datasets (*N* = 8956 proteins). 2211 out of 2865 (77%) proteins identified in the plasma are also identified in the brain. The percentage coverage of proteins in each module is also presented. C) 44 modules of previously generated consensus human brain co-expression network ^11^ visualized in the order of module relatedness following protein re-assignment as described in method. D) Overlap of Hp-enriched plasma proteins (y-axis) with increased abundance in AD depicted in red or decreased abundance in AD shown in blue within brain network modules. The intensity of color shading indicates the degree of overlap. Statistical significance is indicated in the heatmap regions using stars (* *p* < 0.05, ** *p* < 0.01, *** *p* < 0.001). The *p*-values derived from FET were corrected using the BH method. E) The heatmap demonstrates the bicor correlations of each module with CERAD, Braak, and MMSE cognitive scores (* *p* < 0.05, ** *p* < 0.01, *** *p* < 0.001). As mentioned previously, M42 ‘Matrisome’ exhibits the strongest correlation with AD pathology, and several synaptic modules (M1, M4, M5, and M22) display an overall decrease in AD brain. F) Using FET, the cell type nature of each module was assessed by module protein overlap with brain cell-type-specific markers of astrocytes, microglia, neurons, oligodendrocytes and endothelia. The strength of the color shading indicates the degree of cell type enrichment with asterisks denoting statistical significance (* *p* < 0.05, ** *p* < 0.01, *** *p* < 0.001). The FET-derived *p*-values were corrected using the BH method. CTL, control.

### Integration of human brain network modules with heparin-enriched plasma in AD resolves shared pathways reflective of disease pathophysiology

To gain deeper insights into the biological connection between differentially abundant AD plasma proteins and the brain network, we conducted a comprehensive integrated analysis of the two distinct proteomes. Specifically, we examined which brain co-expression modules (**Figure 6C**) exhibited strong overlap with differentially expressed Hp-enriched plasma proteins using a Fisher’s exact test (FET). As depicted in **Figure 6D**, seven brain modules had significant overlap with Hp-enriched proteins increased in AD plasma, while 16 modules exhibited overlaps with proteins significantly decreased in AD plasma. As described previously ^11^, many of these brain modules displayed significant correlations with AD pathology (i.e., CERAD for neuritic plaque burden and Braak staging for the progression of tau neurofibrillary tangles), cognitive status prior to death (MMSE), and brain cell types as illustrated in **Figure 6E-F**. This allowed us to prioritize modules that exhibited significant correlations with AD clinicopathological traits and specific brain cell types and that were also enriched with differentially abundant plasma proteins. We categorized these modules into groups based on their expression trends (**Figure 7**). For example, certain modules displayed concordant changes in AD plasma and brain even in some instances in the asymptomatic phase of disease in the brain. This included modules such as the ‘Matrisome’ (M42), whereas modules associated with ‘Acute-phase response’ (M26) and ‘RNA-splicing’ (M13) were increased in both plasma and brain only in the symptomatic phase of disease. We also observed significant overlap between plasma proteins that are significantly decreased in AD and modules related to synaptic proteins or ER/glycosylation (M1, M8, and M29) that exhibit a gradual reduction in levels within the brain across control, AsymAD, and AD cases. Interestingly, within the modules, the trends were not consistent for all plasma proteins. For example, in M42, while most of the matrisome members were significantly increased in AD brain and plasma, SDC4 and APP exhibited significant decreases in AD plasma, which was in opposition to their changes in the brain. Another class of modules demonstrated divergent abundance trends in AD brain and plasma (M11, M7, M25, M39, M15, and M44). For instance, proteins specifically in M11 ‘Cell adhesion/ECM’, M7 ‘MAPK signaling’ and M25 ‘Sugar metabolism’, such as PTBP1, FHL1 and DCTN2, exhibited increased levels in the AD brain, yet decreased levels in the Hp-enriched AD plasma. Finally, modules that showed significant overlap with the plasma proteome, yet exhibited moderate to low correlations with AD clinicopathological traits are detailed in **Supplemental Figure 5**. These included M40 ‘Antigen binding’ which is highly increased only in the AD plasma, potentially reflecting a peripheral-derived inflammation and immune response to AD pathology in the brain. On the contrary, M37 ‘Endosome’ and M30 ‘Proteasome’ was significantly decreased in AD plasma while not changing in the brain, suggesting a potential failure of protein clearance in AD plasma. In summary, these analyses facilitated the categorization of groups of Hp-enriched plasma proteins based on their associations with AD brain pathophysiology. Additionally, they shed light on the directionality of specific plasma proteins within the network modules. Some of these proteins exhibited consistent increases or decreases in both the brain and plasma, while others displayed divergent changes.

**Figure 7.**
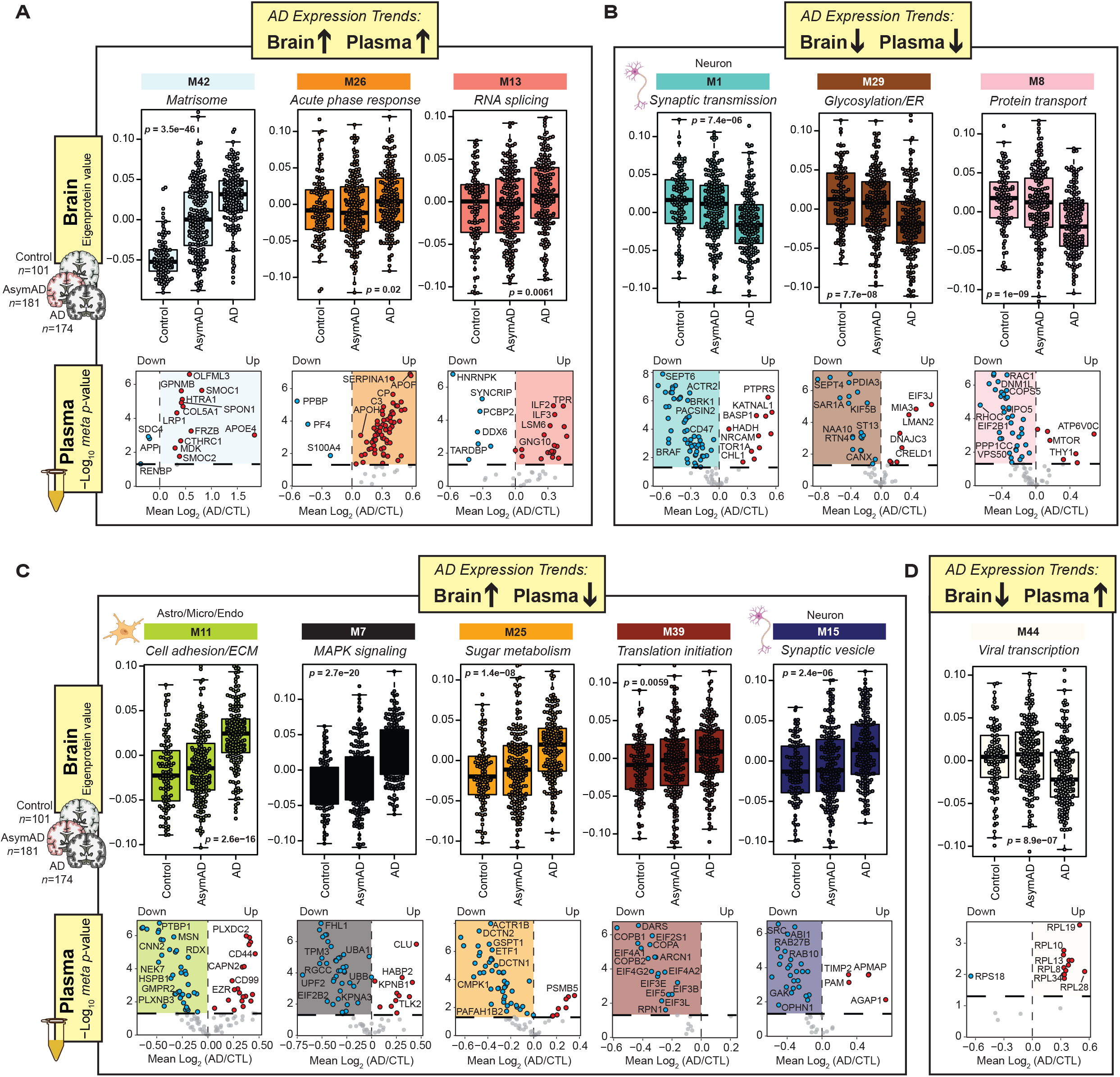
Overlap between human brain network modules associated with AD continuum and differentially abundant heparin-enriched AD plasma proteins. Protein expression trends of the 12 out of 44 (27%) consensus brain network modules ^11^ that exhibited significant correlations with AD clinicopathological traits and were enriched with differentially abundant plasma proteins. Brain module abundance is represented by eigenprotein values of the consensus brain network (control = 101, AsymAD = 181, AD = 174), while volcano plots illustrate the differential abundance (log^2^ fold-change AD vs CTL) of module proteins in the Hp-enriched plasma proteome. The statistical significance of changes in module eigenprotein abundance across the three groups of the consensus brain cohort was assessed using ANOVA with Tukey post-hoc correction. Modules with *p* < 0.05 were considered significant. A) The M42 ‘Matrisome’, M26 ‘Acute-phase response’, and M13 ‘RNA-splicing’ displayed increased protein levels in both the AD brain and plasma. B) The M1 ‘Synaptic transmission’, M29 ‘Glycosylation/ER’ and M8 ‘Protein transport’ showed decreased protein levels in both AD brain and plasma. C-D) The remaining six modules, including M11 ‘Cell adhesion/ECM’, M7 ‘MAPK signaling’, M25 ‘Sugar metabolism’, M39 ‘Translation initiation’, M15 ‘Synaptic vesicle’, and M44 ‘Viral transcription’, exhibited divergent abundance changes in AD brain and plasma. CTL, control; Astro, astrocytes; Micro, microglia; Endo, endothelia.

## Discussion

We utilized heparin affinity chromatography to enrich and improve the detection of matrisome-associated HBPs altered in AD plasma. Using TMT-MS, we identified and quantified nearly 3,000 proteins across two independent sets of plasma samples (*n* = 121) spanning over 10 orders of magnitude in protein concentration in plasma. Despite these plasma samples being processed using a different lot and volume of heparin sepharose beads, on separate MS platforms and nearly a year apart, we observed a high degree of reproducibility in the significance and effect size when comparing differences between AD and controls. Moreover, we present compelling evidence indicating that proteins associated with heparin binding, including those that overlap with the M42 matrisome module in the brain ^11^ are significantly increased in plasma of AD patients. These elevated levels of matrisome proteins in AD plasma also demonstrate a strong correlation with CSF concentrations of Aβ^1-42^, tTau, and pTau181, as well as plasma pTau181 suggesting that these proteins are related to the pathophysiological mechanisms of AD in both the brain and plasma.

One of our main goals was to implement a heparin affinity enrichment for the capture of M42 members, which we have previously shown to be strongly correlated with amyloid and tau pathology in the brain and CSF ^11, 19, 20^. For example, not only are SMOC1 and SPON1 increased in CSF upwards of 30 years in advance of AD ^17^, these proteins are also increased in the preclinical or asymptomatic stage of disease in brain making each high value biomarkers if detected in plasma^11^. Both proteins serve as hub proteins within M42 and maintain their centrality within this module even after analyzing and quantifying the brain proteomic data using an different independent search algorithm. Notably, the new database search output of 456 brain samples across Banner and ROSMAP using FragPipe was able to identify 10% more proteins that were associated with M42. Other newly identified members of M42 include SMOC2 and HGF, which have also been shown to be elevated in AD CSF ^64-66^. Consequently, our findings suggest that both SMOC1 and SMOC2 exhibit elevated levels in the brain and plasma of individuals with AD. SPON1, which was also increased in Hp-enriched plasma in AD, was one of the most consistent M42 proteins measured across proteomic platforms as it was increased in AD by aptamer-based and Olink antibody-based measurements. In total, 16 matrisome proteins overlapping with M42 were differentially abundant in AD plasma following heparin enrichment, where SMOC1, SPON1, OLFML3, GPNMB, and HTRA1 are among the most significant ones elevated. Future studies aiming to measure matrisome proteins in plasma from cohorts like Alzheimer’s Disease Neuroimaging Initiative (ADNI) and the Dominantly Inherited Alzheimer Network (DIAN) will be essential for assessing their prognostic values in predicting disease outcomes.

In our previous consensus analysis of the human brain, we did not incorporate the APOE-specific isoforms (APOE2 and APOE4) into the human protein database ^11^. Since these isoforms differ due to substitutions of cysteine residues with arginine residues, they can be easily distinguished in the human proteome after trypsin cleavage which releases novel peptides, which can then be accurately mapped and quantified using mass spectrometry ^57, 67^. Notably, by integrating genomics and proteomics data from the same individuals, we previously demonstrated that the individuals carrying an APOE ε4 allele exhibited higher M42 levels in brain, and this regulation was not solely driven through the levels of the APOE protein itself ^11^. In this study, the inclusion of the APOE ε4 specific protein isoform rather than APOE in M42 serves to further strengthen the genetic association between APOE ε4 genotype and M42 levels. This also indicates that the APOE ε4 isoform may have a stronger predisposition for interaction with heparin and HBPs within M42 in the brain than other APOE isoforms ^59^. In the future, the implementation of integrated genomics and proteomics pipelines will be needed for assessing whether M42 proteins in plasma are under genetic regulation by APOE4. This will provide valuable insights into whether this relationship is consistent across both the central nervous system and the peripheral system.

Heparin and HS accelerate the formation of Aβ fibrils ^27-29^ and loss of this heparin-APOE binding interaction has been suggested as a possible mechanism for the protection of the *APOE* Christchurch loss-of-function mutation recently described in a *PSEN1* ADAD mutation carrier ^59^. However, more recently, a rare *RELN* variant has been proposed to delay the age of onset of siblings with ADAD who do not carry the Christchurch *APOE* variant ^68^. Like APOE, Reelin (RELN) is also an HBP, but in contrast to the *APOE* Christchurch variant, the RELN variant is associated with heightened interactions with heparin and a consequent reduction in tau phosphorylation via the Dab1 signaling pathway ^68^. It is worth noting that in our study, we observed increased levels of RELN in the Hp-enriched fraction compared to input and FT (**Supplemental Figure 1**). Hence, there appears to be an intricate relationship between heparin interactions, APOE, and AD risk. The exact mechanisms by which HBPs, including APOE and members of M42, influence amyloid deposition and potential clearance still requires further investigation.

Beyond their interactions with APOE and M42 members, heparin and HS alone have been implicated in AD progression and pathogenesis ^69^. Specifically, heparan sulfate proteoglycans (HSPGs) are believed to play a crucial role in mediating the internalization and propagation of specific proteopathic seeds of tau ^70^. Namely, the HSPG glypican-4 (GPC4) has been identified as a contributor to APOE4-induced tau hyperphosphorylation ^71^. It is also noteworthy that 6-O sulfation on HSPGs is presumed to regulate the cell-to-cell propagation of tau ^72^. Interestingly, glypican-5 (GPC5), which is structurally homologous to glypican-4, is a core member of M42 (**Figure 6A**). While it is yet to be established whether glypican-5 plays a role in the regulation of tau internalization, the co-expression between APOE4 and GPC5 in the brain suggests the possibility of such involvement. Evidence supporting the role of HS in the etiology of AD is also emerging in genome-wide association studies (GWAS). For example, GWAS meta-analysis identified the heparan sulfate-glucosamine 3-sulfotransferase 1 gene (*HS3ST1*) as a risk locus associated with late onset AD ^73, 74^. Furthermore, a recent study reported a seven-fold increase in total brain HS in AD compared to controls and other tauopathies ^75^. These findings collectively suggest that dysfunction in HS and HBPs in brain, CSF and plasma may play a central role in the etiology and the clinicopathological presentation of AD.

In addition to the matrisome-associated proteins, we also identified a significant number of other Hp-enriched plasma proteins that exhibited increased levels in AD. Among these proteins were the proteoglycan biglycan (BGN), which is typically found in the extracellular matrix of blood vessels ^76, 77^, and Endothelial Cell–Specific Molecule 1 (ESM1), also known as endocan. Both proteins play roles in regulating endothelial cell function and angiogenesis, and they have been implicated in processes related to inflammation and vascular disease ^76-78^. Furthermore, we observed an elevation in Colony-Stimulating Factor 1 (CSF1) in the plasma of individuals with AD. CSF1 primarily functions in the regulation of the immune system ^79^. Elevated levels of CSF1 in plasma have been associated with various diseases and conditions, including inflammation, cancer, and certain autoimmune disorders ^80-82^. Taken together, these findings suggest that widespread systemic inflammation may also manifest in the plasma of individuals with AD. However, whether this phenomenon is specific to AD or extends to other subtypes of dementia remains a subject that requires further investigation.

In this study we also leveraged the consensus brain protein co-expression network to explore the relationship between the Hp-enriched plasma and brain proteomes. Within the network modules, certain plasma proteins consistently exhibited increases or decreases in AD that mirrored changes in the brain, while others displayed divergent alterations as previously described ^31^. Among those consistently increased, beyond M42 members, were proteins mapping to M26 in the brain, associated with the ‘Acute phase response’, which suggests that proteins related to complement activation potentially associated with immune function are enriched in AD plasma. Notably, proteins of interest in M26 included SERPINA3, which was recently identified through a large-scale analysis of the plasma proteome using Mendelian randomization as potentially causal in AD pathogenesis ^83^. Additionally, we observed proteins in plasma that map to neuronal modules related to synaptic biology in the brain which displayed consistent decreases in AD. Whether this change in plasma reflects synapse loss in the brain will require further investigation. Nevertheless, it is intriguing that a signature of synaptic loss typically associated with cognitive decline in brain and CSF appears in the Hp-enriched plasma in AD. There was also some discordance in the direction of change between AD plasma and brain proteomes. For instance, proteins specifically in M7 ‘MAPK signaling’ and M25 ‘Sugar metabolism’, exhibited increased levels in the brain, yet decreased levels in the Hp-enriched plasma. This trend contrasts with the direction of change observed in CSF ^31^, where glycolytic signature is increased in AD even in the preclinical phase ^14, 31, 84^. The precise mechanisms underlying this discordance remain unclear. However, it is plausible that blood-brain barrier dysfunction might contribute to these differences ^85^, where proteins increased in plasma are decreased in CSF, and vice versa.

Although our goal was to capture HBPs in plasma, a consequence of the heparin affinity enrichment was the clearance of highly abundant proteins such as albumin. This reduction in albumin from the Hp-enriched fraction resulted in the comprehensive coverage of nearly 3,000 plasma proteins following high-pH off-line fractionation and TMT-MS. Thus, in contrast to immune-depletion methods ^86^, antibody-free affinity enrichment-based approaches utilizing nanoparticle, cationic/anionic or hydrophobic/hydrophilic-based strategies appear to substantially enhance plasma proteome coverage through MS-based technologies ^87^. Collectively, this progress marks a step toward overcoming one of the major limitations of plasma MS-abased proteomics, which is the vast dynamic range of protein abundances. Furthermore, the utilization of more advanced mass spectrometers, such as the Orbitrap Astral, which quantifies five times more peptides per unit time than state-of-the-art Orbitrap mass spectrometers ^88^, is expected to significantly enhance the depth of proteome coverage in plasma when employing these affinity enrichment strategies. This is of particular significance due to the complementary coverage of the Hp-enriched plasma proteome by TMT-MS in contrast to the SomaScan and Olink platforms, which will further enhance the depth of the plasma proteome when measurements are integrated across platforms ^31^.

While this study provided a comprehensive proteomic analysis of Hp-enriched plasma from human subjects, several limitations should be acknowledged. Notably, the study participants predominantly consisted of non-Latino white individuals. Recent reports have highlighted disparities in AD prevalence, with Black and Hispanic populations showing a higher likelihood of developing AD compared to older white Americans ^89-91^. Additionally, it has been observed that cognitively impaired African American individuals have lower levels of CSF tTau and pTau compared to Caucasians ^40^. An important ongoing initiative of the Accelerating Medicines partnership for AD (AMP-AD) ^92^ is the inclusion of African American and Hispanic individuals in plasma biomarker studies. Research efforts employing heparin enrichment techniques should aim to encompass a more diverse participant population to better capture the complexities of AD across different racial and ethnic groups. Future studies that investigate the interplay between age, sex, and race within the Hp-enriched plasma proteome will yield valuable insights. Moreover, our plasma proteomics study exclusively focused on control and symptomatic individuals who were AD biomarker positive based on their tau/amyloid ratio in the CSF. Future plasma proteomic studies aimed at exploring the pre-symptomatic stages of AD before cognitive impairment manifests will be needed to identify Hp-enriched plasma proteins that undergo early changes in the disease course. Nevertheless, this study offers a global view into the Hp-enriched plasma proteome, reinforcing a hypothesis that increased matrisome proteins are shared between the brain and blood in AD.

## Acknowledgements

We thank the Seyfried lab and members of the Emory Center for Neurodegenerative Disease (CND) for helpful comments and discussions.

## Author contributions

Conceptualization, D.M.D, E.J.F. and N.T.S.; Methodology, Q.G., L.P., Y.L., K.X., D.M.D. B.R.R., J.J.L and N.T.S; Investigation, Q.G., L.P., E.B.D., J.J.L., A.I.L., and N.T.S.; Formal Analysis, Q.G., E.B.D., A.S. ; Writing – Original Draft, Q.G., L.P., D.M.D., and N.T.S.; Writing – Review & Editing, Q.G., L.P., E.J.F. D.M.D., E.B.D, E.C.B.J., J.J.L., A.I.L., and N.T.S.; Funding Acquisition, J.J.L, A.I.L. and N.T.S.; Resources, J.J.L. A.I.L. ; Supervision, A.I.L., and N.T.S. All authors read and approved the final manuscript.

## Availability of data and materials

Raw mass spectrometry data and pre- and post-processed plasma protein abundance data and case traits related to this manuscript are available at https://www.synapse.org/HeparinPlasma. The results published here are in whole or in part based on data obtained from the AMP-AD Knowledge Portal (https://adknowledgeportal.synapse.org). The AMP-AD Knowledge Portal is a platform for accessing data, analyses and tools generated by the AMP-AD Target Discovery Program and other programs supported by the National Institute on Aging to enable open-science practices and accelerate translational learning. The data, analyses and tools are shared early in the research cycle without a publication embargo on secondary use. Data are available for general research use according to the following requirements for data access and data attribution (https://adknowledgeportal.synapse.org/#/DataAccess/Instructions).

## Abbreviations

HBP: Heparin binding protein
AD: Alzheimer’s disease
CSF: cerebrospinal fluid
Aβ: β-amyloid
MS: mass spectrometry
APOE: apolipoprotein E
APP: amyloid precursor protein
APOE4: isoform E4 of APOE
APOE2: isoform E2 of APOE
Hp-enriched: heparin-enriched
tTau: total tau
pTau: phosphorylated tau
HS: heparan sulfate
ECM: extracellular matrix
PET: positron emission tomography
TMT- MS: tandem ass tag mass spectrometry
LFQ: label-free quantification
CERAD: Consortium to Establish a Registry for Alzheimer’s Disease
MMSE: Mini-Mental State Examination
DP: diluted plasma
Hp- depleted: heparin-depleted
FT: flowthrough
PD: Proteome Discoverer
FP: FragPipe
FDR: false discovery rate
GIS: global internal standard
DDA: data-dependent acquisition
AGC: automatic gain control
HCD: higher-energy collision-induced dissociation
GO: gene ontology
bicor: biweight midcorrelation

## Supplemental Figure Legends

**Supplemental Figure 1.**
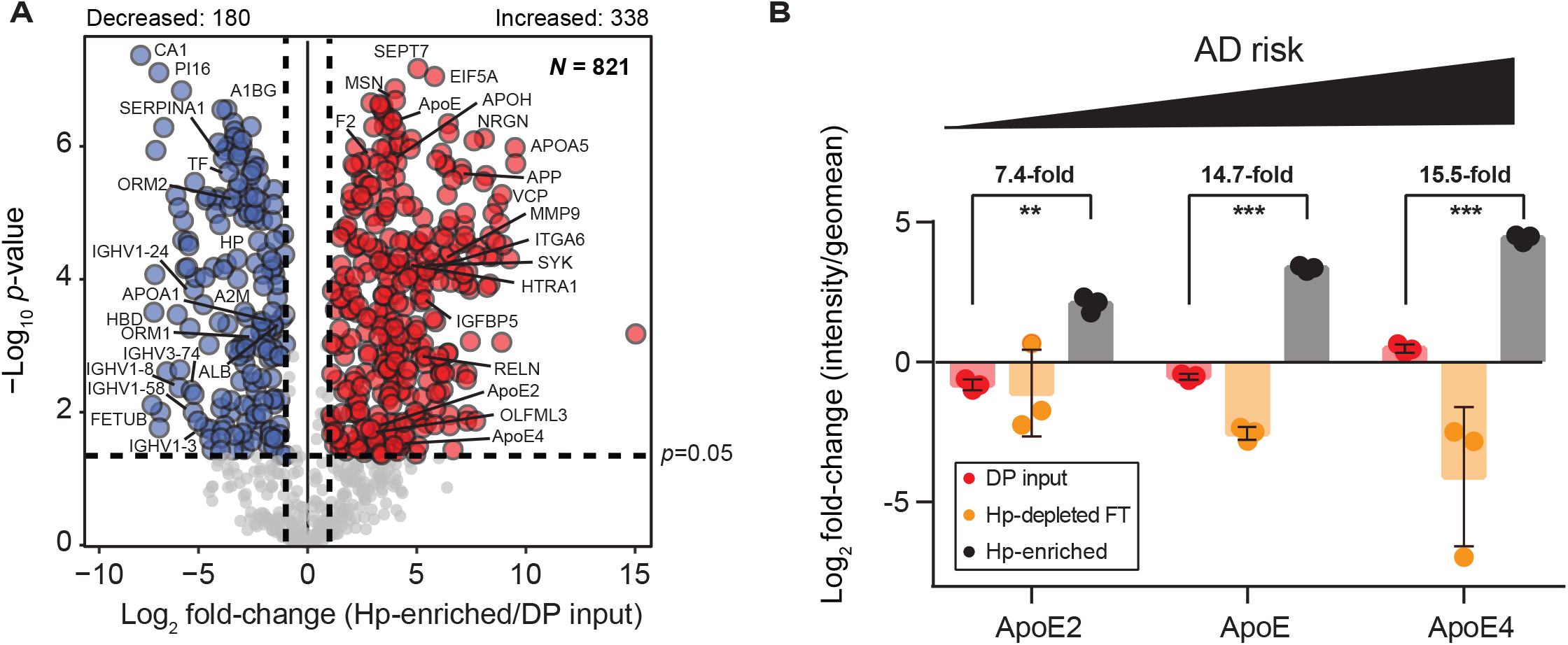
Comparing heparin-enrichment across APOE isoforms. **A)** The volcano plot shows differentially enriched proteins in the Hp-enriched fractions (*n* = 3) of pooled plasma sample compared to the DP inputs (*n* = 3). Red circles represent significant Hp-enriched proteins in plasma and examples of AD-related HBPs are highlighted, in addition to APOE isoforms. Blue symbols represent proteins significantly depleted from the Hp-enriched fractions. The significance cutoff is *p* < 0.05 (ANOVA with Tukey post-hoc correction) and fold-change >2. **B)** The bar plot shows abundance changes of APOE2, APOE and APOE4 isoforms across three fractions of pooled plasma (DP input = 3, Hp-depleted FT = 3, Hp-enriched = 3). The y-axis is calculated by log^2^ fold-change of protein intensity over their geomean across all 9 samples. The significance of difference was determined by ANOVA with Tukey post-hoc correction and denoted with stars (* *p* < 0.05, ** *p* < 0.01, *** *p* < 0.001).

**Supplemental Figure 2.**
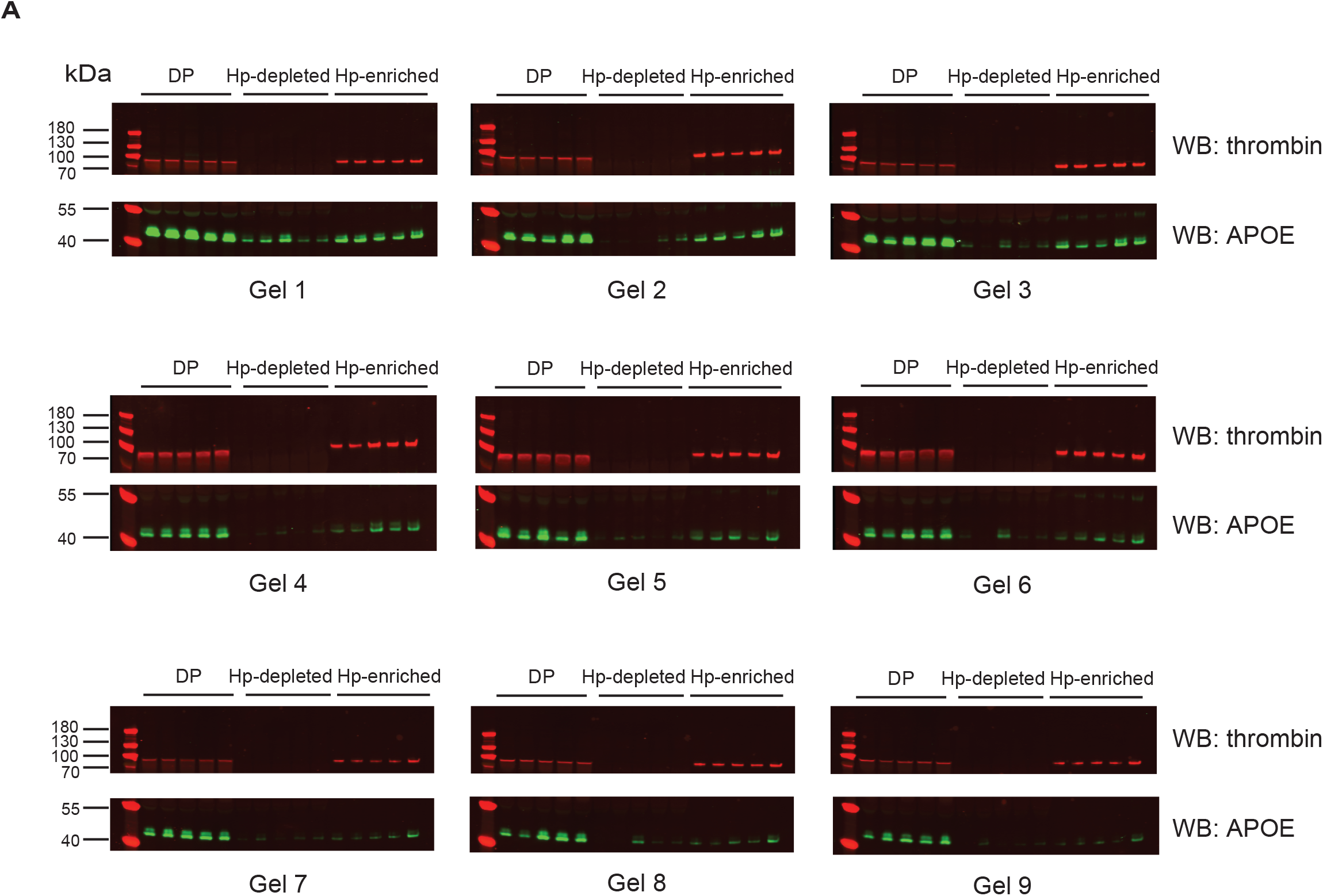
Western blotting of thrombin and APOE in Set 1 (*n* = 36). A) Western blotting was conducted on DP input, Hp-depleted FT, and Hp-enriched fraction obtained from Set 1 samples, which included 18 control and 18 AD cases. Thrombin and APOE were the target HBPs of interest. On each gel, 2 control samples, 2 AD samples, and 1 GPS sample (see method) were loaded for each fraction. The results show that both thrombin and APOE were depleted from the Hp-depleted FT and enriched in the Hp-enriched fraction. WB: western blotting.

**Supplemental Figure 3.**
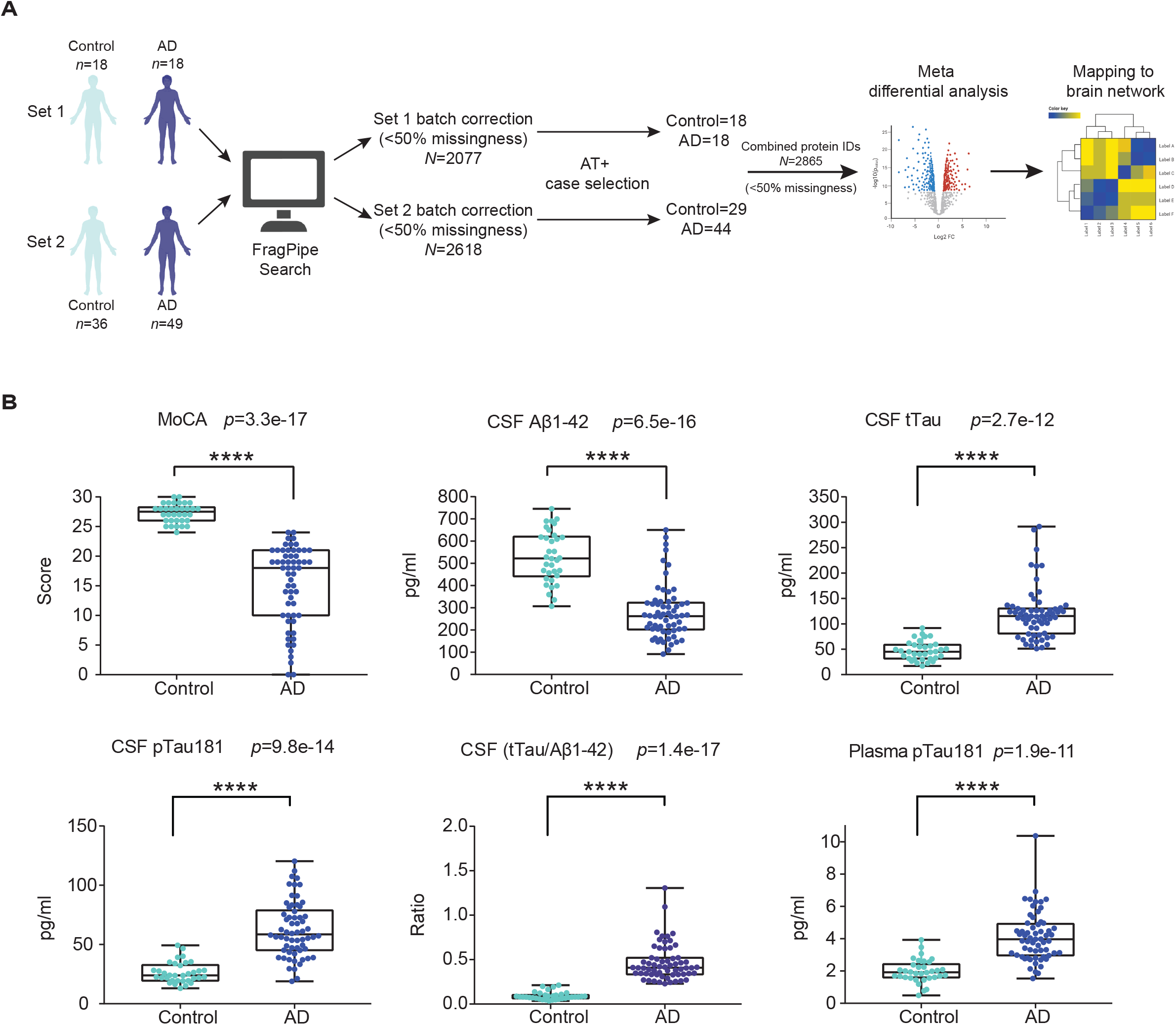
Workflow of the data analysis pipeline across two Hp-enriched plasma datasets. **A)** Set 1 and Set 2 were jointly analyzed using FP, followed by independent batch-specific variance correction procedures, which included TAMPOR and batch-regression. Subsequently, 12 cases from Set 2 that did not meet the AT+ threshold criteria were excluded. A total of 109 samples and 2865 total proteins were selected for further analysis, with 13 overlapping control samples between the two datasets. B) Measurements of various AD biomarkers, including cognition (MoCA score), CSF Aβ^1-42^, CSF tTau, CSF pTau181, CSF ratio of tTau/Aβ^1-42^, and plasma pTau181, were obtained for the selected unique cases (*n* = 96). Significance levels determined by Student’s *t*-test are denoted by overlain asterisks; \**p* < 0.05, \*\**p* < 0.01, \*\*\**p* < 0.001, \*\*\*\**p* < 0.0001.

**Supplemental Figure 4.**
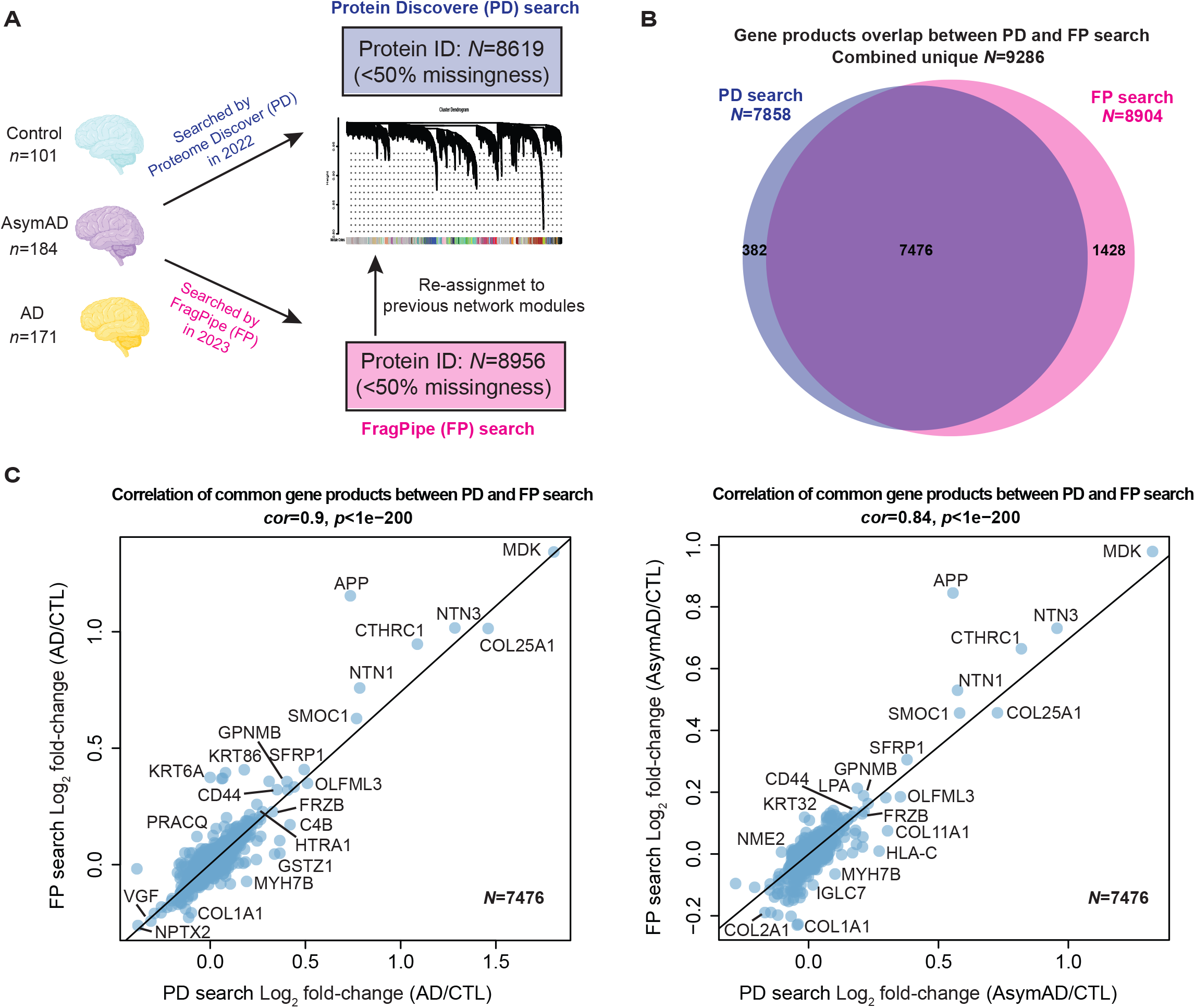
Comparing TMT-MS proteomic measurements of human brain generated by FP and PD. A) 456 raw files collected from the ROSMAP and Banner cohorts as previously described ^11^ underwent a database search using FP (see method), resulting in the identification of 8956 UniprotID-identified proteins, each with measurements available in 50% or more across 456 individual cases (control = 101, AsymAD = 181, AD = 174). These proteins were subsequently assigned to one of the 44 consensus network modules ^11^ by re-calculating the kME (bicor correlation to module eigenprotein) for each protein and assigning it to the module that exhibited the highest correlation ^53^. B) The number and overlap of unique gene products identified in the FP and the PD outputs, with 7476 overlapping between the two datasets. FP provided an additional 1428 unique gene products compared to PD, resulting in an 18% increase in proteome coverage. C) Scatter plots illustrate the correlation between log^2^ fold-change for AD vs CTL (left, *cor* = 0.9, *p* < 1e^-200^) and AsymAD vs CTL (right, *cor* = 0.84, *p* < 1e^-200^), using common gene products (*N* = 7467) found in both FP and PD search results. CTL, control; *cor*, Pearson correlation of coefficient.

**Supplemental Figure 5.**
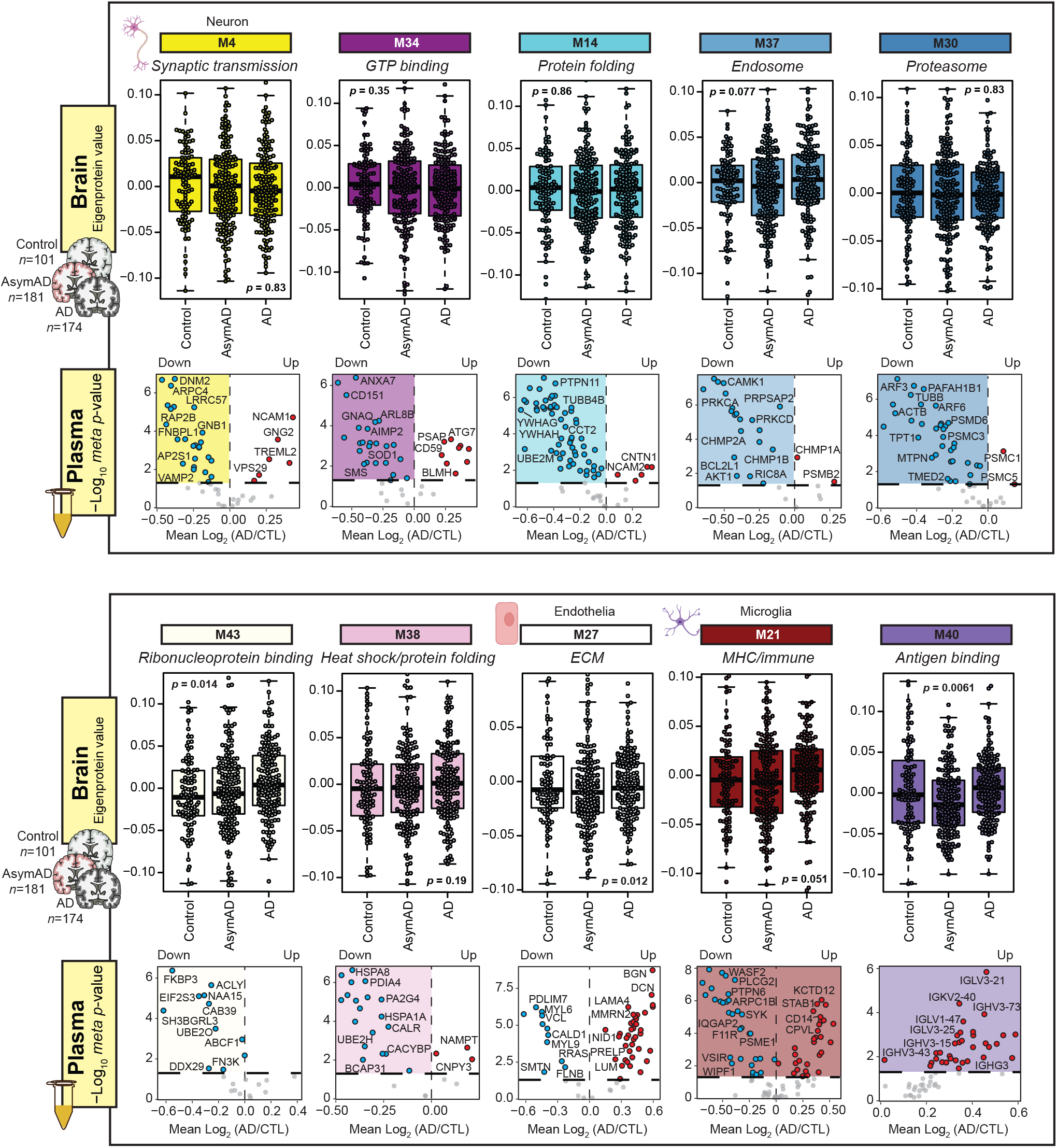
Overlap between human brain network modules with low correlation to AD pathology and differentially abundant Hp-enriched plasma proteins in AD. Protein expression trends are examined for the 10 modules that exhibit significant overlap with differentially abundant Hp-enriched plasma proteome but demonstrate moderate to low correlation with AD clinicopathological traits in the brain. Brain module abundance is quantified by eigenprotein values derived from the consensus brain dataset ^11^ (control = 101, AsymAD = 181, AD = 174), while volcano plots illustrate the differential expression (log^2^ AD vs CTL) of module proteins in the Hp-enriched plasma proteome. The statistical significance of changes in module eigenprotein abundance across the three groups in the consensus brain cohort was assessed using ANOVA with Tukey post-hoc correction. Modules with *p* < 0.05 were considered significant. Among these modules, M4 ‘Synaptic transmission’, M34 ‘GTP binding’, M14 ‘Protein folding’, M37 ‘Endosome’, M30 ‘Proteasome’, M43 ‘Ribonucleoprotein binding’ and M38 ‘Heat shock/protein folding’ consist of proteins with decreased abundance in AD plasma, while M40 ‘Antigen binding’ exclusively contains increased plasma proteins. M27 ‘ECM’ and M21 ‘MHC/immune’ exhibit a balanced representation of both increased and decreased plasma proteins. CTL, control.

